# “Comparative genomics and pangenomics of vancomycin resistant and susceptible *Enterococcus faecium* from Irish hospitals across 20 years”

**DOI:** 10.1101/2021.11.22.469549

**Authors:** R.J. Leigh, C. McKenna, R. McWade, B. Lynch, F. Walsh

**Affiliations:** Department of Biology, Maynooth University; Department of Microbiology, Mater Misericordiae University Hospital

**Keywords:** Vancomycin resistance, Enterococcus, Evolution, Ireland, CC17

## Abstract

*Enterococcus faecium* has emerged as an important nosocomial pathogen, which is increasingly difficult to treat due to the genetic acquisition of vancomycin resistance. Ireland exhibits a recalcitrant vancomycin resistant bloodstream infection rate compared to other developed countries. A set of 28 vancomycin resistant isolates was sequenced to construct a dataset alongside 61 other publicly available Irish genomes. This dataset was extensively analysed using *in-silico* methodologies and uncovered distinct evolutionary, coevolutionary, and clinically relevant population trends. These results suggest that a stable (in terms of genome size, GC%, and number of genes), yet genetically diverse population (in terms of gene content) of *Enterococcus faecium* persist in Ireland with acquired resistance arising *via* plasmid acquisition (*vanA*) or to a lesser extent, chromosomal recombination (*vanB*). Population analysis described five clusters with one cluster partitioned into four clades which transcend isolation dates. Pangenomic and recombination analyses revealed an open (whole genome and chromosomal specific) pangenome illustrating a rampant evolutionary pattern. Comparative resistomics and virulomics uncovered distinct chromosomal and mobilomal propensity for multidrug resistance, widespread chromosomal point-mutation mediated resistance, and chromosomal harboured arsenals of virulence factors. Comparative phagomics revealed a core prophagome of three prophages throughout the dataset. Interestingly, a potential difference in biofilm formation strategies was highlighted by coevolutionary analysis, suggesting differential biofilm genotypes between *vanA* and *vanB* isolates. These results highlight the evolutionary history of Irish *Enterococcus faecium* isolates and may provide an insight into underlying infection dynamics in a clinical setting.

## Introduction

The genus *Enterococcus* (Firmicutes; Bacilli; Lactobacilli; Enterococcaceae) are commonly observed in diverse biomes such as soil, surface water, wastewater, and as commensal inhabitants of the higher chordate (inclusive of human) gastrointestinal (GI) tract, vaginal tract, and epidermis (1, 2). Two species, *Enterococcus faecium* and *Enterococcus faecalis*, are aetiological of a cohort of moderate to severe conditions when migrated from the gastrointestinal tract, especially in the immunocompromised or convalescing host (3, 4). Both *E. faecium* and *E. faecalis* migration can lead to caries, cellulitis, cholecystitis, cystitis, endocarditis, endodontits, periodontitis, peri-implantitis, postoperational peritonitis, sepsis, and neonatal meningitis (5–8). The clinical significance of these species was amplified as antimicrobial resistance (AMR) began to evolve and disseminate throughout diverse environments (2, 9).

Vancomycin is the drug-of-last-resort for recalcitrant infections with diverse resistance profiles (10), inclusive of the prominent nosocomial pathogen, methicillin-resistant *Staphylococcus aureus* (MRSA) and *E. faecium* (11, 12). In recent years, vancomycin resistance has been observed in a number of human pathogens, often resulting in clinical complication and extended hospital stay (6,13–16). Ireland has one of the highest vancomycin resistant *E. faecium* (VRE) rates in Europe where 38.4% of blood stream infection isolates displaying resistance (ECDC: https://atlas.ecdc.europa.eu/). In particular, nosocomial VRE infection are of concern due to the rapid dissemination throughout this environment (4,15,17). Vancomycin resistance genes have been detected both on the chromosome and mobile elements (mobilome) and often accompanies other resistance genotypes (6,18–20). Commonly observed *vanA* mobilome co-resistance phenotypes are observed towards tetracycline, erythromycin, and aminoglycosides (21–24). Additionally, *E. faecium* almost ubiquitously displays chromosomal-mediated resistance to aminoglycosides (*via AAC*(6’)-*Ii*), macrolides (*via efmA*, *efrAB* and *msrC*), and fluoroquinolones (*via gyrA* and *parC* mutations or upregulation of efflux), rifamycin (*via efrAB*), and to clindamycin, quinupristin-dalfopristin, and dalfopristin (*via lsaA*) (6,9,24–26).

In many countries, including the Republic of Ireland, vancomycin resistance is prominently derived from plasmid mediated *vanA*, and less commonly from chromosomal mediated *vanB* (2,27,28). Both *vanA* and *vanB* are D-Ala-D-Ala ligases (EC: 6.3.2.4) and confer resistance by constructing D-Ala-D-Lac as an alternative substrate during peptidoglycan synthesis, thus reducing vancomycin-peptidoglycan binding affinity (29–31). Aside from their associated genomic location (mobilomal *vs* chromosomal), *vanA* also confers resistance to teicoplanin (another glycopeptide antibiotic) whereas *vanB* has not been observed to induce this effect (32–34).

Metal and biocide resistance is of growing concern in a plethora of environments, and especially in clinical settings (35). Correlations have been reported between metal resistance and drug resistance in pathogenic bacteria, suggesting a potential co-evolutionary pressure on both mechanisms (36–38). A range of metals are employed in healthcare for their intrinsic antimicrobial properties, for example silver embedded plasters and copper plated door handles (39–41). Previous studies have reported chromosomal mediated resistance to copper, silver, selenium, and hydrogen peroxide in *E. faecium*, suggesting a growing resistance to passive protection strategies (42–44).

Further to their resistance mechanisms, *Enterococcus* spp. employ a small cohort of effective virulence factors during pathogenesis, allowing for adhesion, biofilm formation, invasion, immunomodulation, and the synthesis of secreted toxins, enzymes, and peptides (such as bacteriocins) into local environments to inhibit competition (9,45,46). Biofilm formation is of critical importance in vancomycin resistant *Enterococcus* (VRE) infections, due to difficulty of clearance and reduced antimicrobial penetration rates (42, 47).

When isolated from human samples, the average *E. faecium* genome contains 2,765 ± 187 genes, yet the pangenome contained 12,457 when sampled from 161 genomes (26). These results, and other pangenomic analyses (*e.g*. (48)) illustrate the genomic plasticity and evolutionary capacity of *E. faecium*. While a major proportion of this variance is attributed to mobile genetic elements (49), to our knowledge, no study has been conducted on chromosomal pangenomic variance, so the extent of mobilome enticed variance is not fully elucidated. The integration of phages (prophages) into bacterial chromosomes can introduce genetic novelty (50–52). While a wide array of *Enterococcus* phages have been identified (53, 54), and parasite-host co-evolutionary trajectories have been explored (55), phage impact on symbiotic co-evolutionary trajectory has yet, to our knowledge, been explored.

The aims of this study were to compare the pan-genomes, mobilomes and chromosomes of the available genome sequences of vancomycin resistant or susceptible *E. faecium* across the timeframe that VRE (analysed by the ECDC: 2002 to 2019) increased in prevalence from 11.1% to 38.4% in Ireland (available at https://atlas.ecdc.europa.eu/public/index.aspx?Dataset=27&HealthTopic=4). Specifically, this study provides an insight into the pangenomics of *E. faecium* from three studies in Republic of Ireland using a comparative genomic, resistomic, and virulomic lens. It is now possible to derive intrapangenome correlations (Whelan *et al.*, 2020) so these approaches shall be explored to provide additional insight.

A recent study of the global dissemination of *E. faecium* identified that it has two main modes of genomic evolution: the acquisition and loss of genes, including antimicrobial resistance genes, through mobile genetic elements including plasmids, and homologous recombination of the core genome (49). Unfortunately, there were no Irish isolates contained within this study.

## Methods

### Microbiological analysis

Twenty-eight isolates identified as vancomycin resistant *Enterococcus faecium* during 2018 and 2019 were collected by the *Mater Misericordiae* University Hospital (MMUH) in Dublin. The sample metadata are described in SI Table 1. Antimicrobial susceptibility testing was performed at the MMUH according to EUCAST guidelines (https://www.eucast.org/ast_of_bacteria/) and subsequently at Maynooth University using disk testing according to the CLSI guidelines. The isolates were investigated for resistance to the following antibiotics: (ciprofloxacin, erythromycin, chloramphenicol, vancomycin, tetracycline, ampicillin, and linezolid) (SI Table 2).

### DNA extractions genome sequencing

DNA was extracted from each of the 28 isolates using the Macherey-Nagel Nucleospin microbial DNA isolation kit according to the manufacturer’s instructions. The extracted DNA was sequenced by Novogene using the “Bacterial resequencing” service on the Illumina with PE150 and Q30 ≥ 80%. This provided >100X coverage of each genome.

### Genome assembly

The 28 samples sequenced for this study and all reads associated with Irish VRE genomes from British Society for Antimicrobial Chemotherapy study PRJEB4344 (hereafter referred to as the “BSAC” isolates (57)) were downloaded from the NCBI sequence read archive (SRA; (58)). Each read pair (sample) was subjected to adapter removal and quality trimming using TrimGalore *v.*0.6.6. (59) using default settings. Adapter removal during the TrimGalore pipeline was powered by CutAdapt *v.*3.0 (60) and FastQC *v.* 0.11.9 (61). Each sample was assembled using Unicycler *v.*0.4.7 (62) using default paired-end settings. Unicycler utilised SPAdes *v.*3.14.1. (63) to assemble reads and used Bowtie2 *v.*2.4.2. (64), Pilon *v.*1.23 (65), BLAST *v.*2.11 (66, 67), and samtools *v.*1.11 (68) to further complete the assembly. Each assembly was quality assessed using CheckM (69) using the *Enterococcus faecium* database (SI Table 3) and sequence typed using MLST *v.*2.19.0 (Jolley and Maiden, 2010; Seemann, 2014) using default settings (SI Table 4; Figure 1). Instances where a ST could not be completely identified, the approximated alleles were used to approximate a ST using the “search by locus combinations” option for *E. faecium* using PubMLST (www.pubmlst.org (72)). Each assembly was separated into “chromosomes” and “plasmids” (hereafter referred to as “mobilomes” as complete plasmids were not always guaranteed and were treated collectively as an extrachromosomal entity) using Platon *v*.1.5.0 (73). These partitioned assemblies were analysed alongside their concatenated “whole genome” assemblies. Assemblies that had a reported completeness percentage ≤ 95% were retained for further analyses.

**Figure 1:**
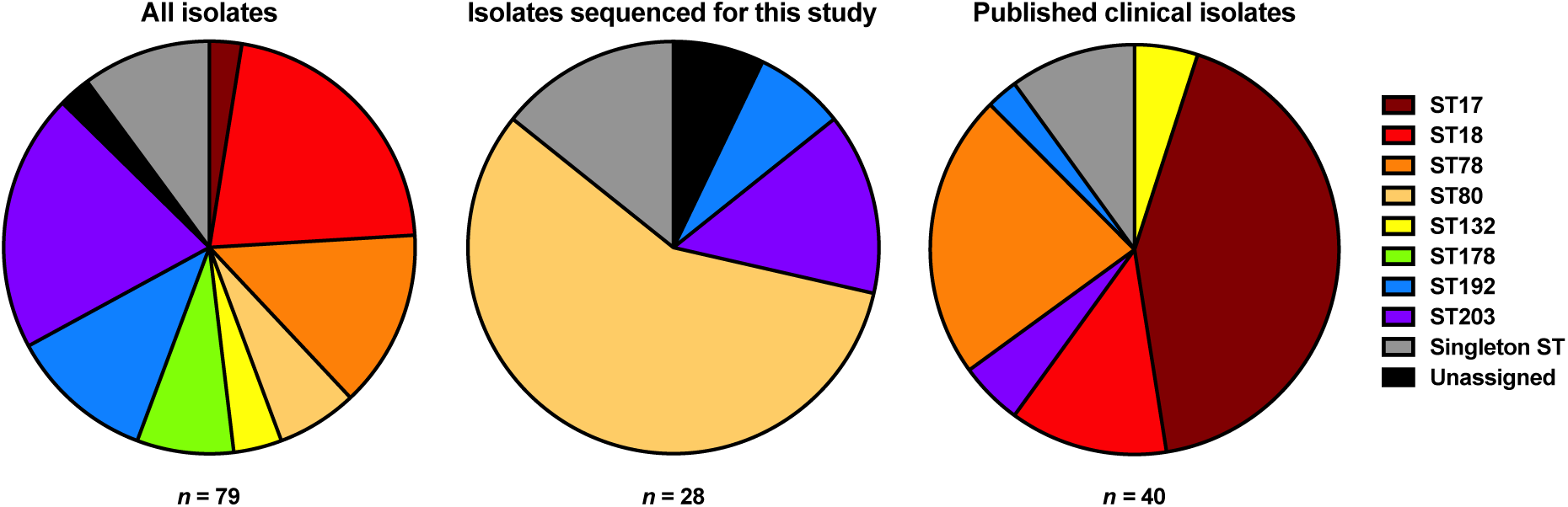
Distribution of sequence types between different studies

### Additional genomes

A total of 11 *Enterococcus faecium* genome assemblies were available on NCBI assembly and attributed to Ireland in their respective metadata (from study PRJNA521309; (74)). As above, these assemblies were quality checked using CheckM, sequence typed using MLST, and separated into chromosomal and mobilomal components using Platon. These 11 isolates were all sampled in county Cork and shall be hereafter referred to as the “Cork” isolates.

### Plasmid containment analysis

As plasmids were contiguous, a containment analysis was used to determine the closest relatives of the mobilome. Each containment analysis was performed using the “screen” algorithm in MASH *v.* 2.2.2. (75) against PLSDB (76) with a *P-*value stringency cut-off of 0.1 and an identity stringency cut-off of 0.99 to replicate the parameters used during the construction of PLSDB (Table SI 5).

### Genome annotation

Each assembly dataset (whole genome, chromosomal, and mobilome) was annotated using Prokka *v.*1.14.6 (Seemann, 2014) using default settings. Prokka used Prodigal *v.*2.6.3. (78) to predict gene sequences, Aragorn *v*.1.2.38 (79) to detect tRNA sequences, and Minced *v*.0.4.0 (80) to detect CRISPR sequences, SignalP *v.*4.0 (81) to detect signal peptides, and HMMER *v.*3.3.1 for protein similarity searching (82), BioPerl *v.*1.7.2 for file manipulation (83), and barrnap *v.*0.9 for rRNA profiling (84). Each protein sequence in each assembly dataset was further annotated using InterProScan *v*.5.45-80.0 (85) using the “--appl PfamA” and “--goterms” to assign Pfam domains (86) and Gene Ontology terms (87) respectively. Finally, each assembly dataset was searched for secondary metabolite biosynthesis genes using a set of tools with their respective default settings (unless otherwise stated below): GECCO *v.*0.6.3 (88) and AntiSMASH *v*.5 (89), for transposable elements using MobileElementFinder *v*.1.0.3. (90), for antimicrobial resistance using ABRicate *v.*1.0.1 (Seemann, 2014) with the associated CARD database (92) and PointFinder *v.*1 (using the *Enterococcus faecium* database (93)), for virulence factors using Abricate with the associated VFDB dataset (94), and for metal (and biocide) resistance using BacMet *v.*2.0 (95). As BacMet is published as an amino acid dataset, it was first backtranslated to a representative nucleotide sequence (using translation table 11) using the “backtranseq” function in EMBOSS *v.*6.6.0.0 (96) with codon usage tables for *Enterococcus faecium* (the subject of this study); *Enterococcus faecalis*, *Staphylococcus aureus*, *Klebsiella pneumoniae*, *Acinetobacter baumanii*, *Pseudomonas aeruginosa*, *Enterobacter* spp. (ESKAPE pathogens); *Escherichia coli* (model organism), *Treponema pallidium*, *Neisseria gonorrhoeae, Chlamydia trachomatis* (prevalent bacterial STIs), *Clostridium botulinum, Campylobacter jejuni, Listeria monocytogenes,* and *Vibrio parahaemolyticus* (prominent foodbourne pathogens), *Helicobacter pylori* (common gastric pathogen) and *Clostridiodes difficle, Legionella pneumophilia,* and *Mycobacterium tuberculosis* (common nosocomial infections). Codon usage tables were obtained from the Kazusa genome research institute (https://www.kazusa.or.jp/en/). To mitigate false negatives, ABRicate was ran with a 50% minimum percentage identity stringency score (as opposed to the default 80%) to allow for the detection of full-length homologs that may have been otherwise undetected (Collated results for AntiSMASH and ABRicate are given in SI Tables 6-10).

### Genome characteristic statistical analysis

The sum of coding genes, genome size (Mbp), genome density (the mean number of genes per Mbp), and guanine-cytosine content (GC%) was calculated for the chromosomal and mobilomal datasets for each sample (SI Table 11; Figure 2). Summary statistics (mean, median, standard deviation, and variance) was computed for each data series (SI Table 12). Two-tailed Welch’s *t*-tests (H_0_:μ*_a_*=μ*_b_*;H_A_:μ*_a_*≠μ*_b_*; (97, 98)) were used to compare genome density and GC% between chromosomal and mobilomal datasets, a Bonferroni-Dunn correction (*P*_BD_ = *P* × *n*_comparisons_; *n*_comparisons_ *=* 2; (Bonferroni, 1936; Dunn, 1961)) was used to control Type-I errors and instances where *P*_BD_ ≤ 0.005 were considered statistically significant. A *P* ≤ 0.005 will be used to determine significance in all pairwise test comparisons to control for potential Type I and Type II errors (101). Summary statistics were calculated for gene length in each genomic subset (chromosomal and mobilomal), distributions were assessed for normality using a Kolmogorov-Smirnoff test (H_0_:∼*X*=*N*(μ,σ);H_A_:∼*X*≠*N*(μ,σ); (102, 103)) and equivariance using a Levene’s test (H_0_:σ^2^ =σ^2^ ;H_A_:σ^2^ ≠σ^2^ ; (Levene, 1960)). All samples were observed to be non-normally distributed and 26 of 79 were observed to have unequal variances (Levene’s test *P* ≤ 0.05). Considering these trends, a Brunner-Munzel test (H_0_:*B*=0.5;H_A_:*B*≠0.5) was used to determine whether chromosomes were statistically more likely to have longer genes than mobilomes. A Bonferroni-Dunn correction was applied (*n*_comparisons_ = 79) and instances where *P_BD_* ≤ 0.005 were considered statistically significant (SI Table 12).

**Figure 2:**
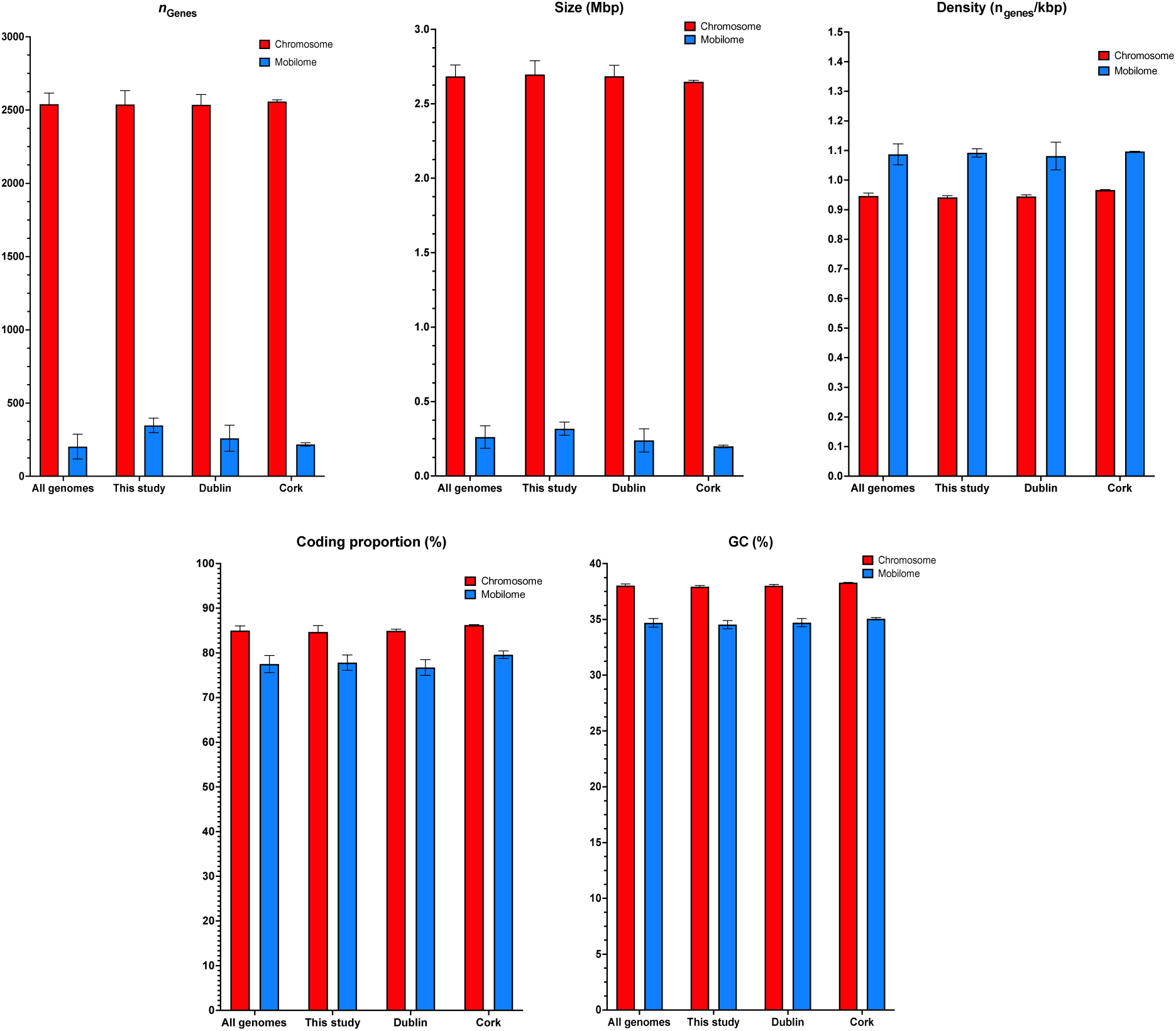
Comparison of different genome characteristics between chromosomes and plasmids across different studies.

### Genome relatedness

Chromosomal sequences from each isolate were all-*vs*-all compared using MASH *v.* 2.2.2. (75) and instances where the reported distance (*D*) ≤ 0.05 with a *P*-value ≤ 0.05 extracted as an edge list and visualised as a network with Gephi *v.*0.9.2 (104) using the Fruchterman-Reingold algorithm (105) (Figure 3). This procedure was repeated for mobilomal sequences (Figure 4).

**Figure 3:**
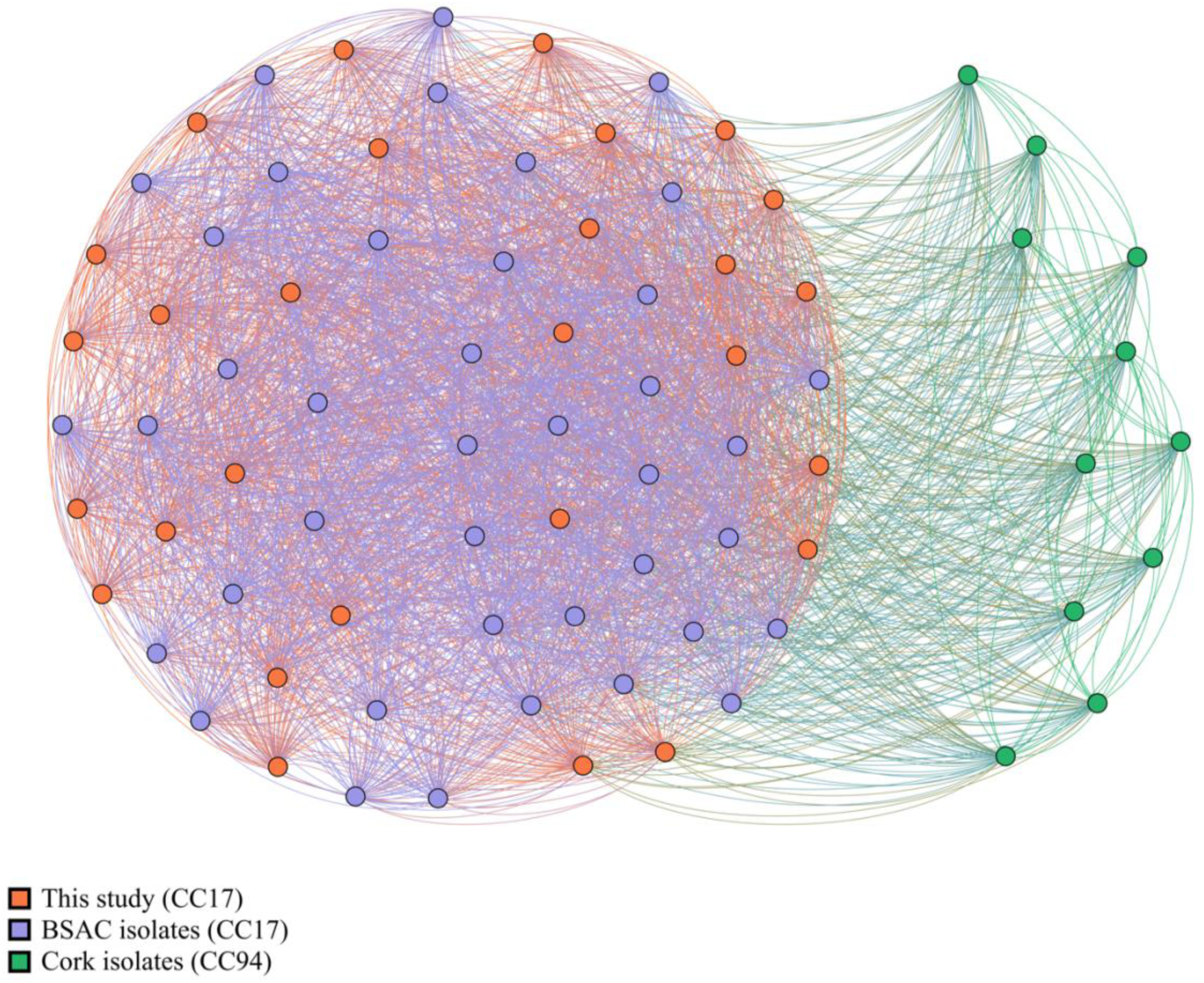
Genomic distance cluster analysis of chromosomal sequences from all isolates. All CC17 isolates cluster together and all CC94 isolates cluster together (whereby all isolates share an edge). Approximately half of CC17 cluster with CC94.

**Figure 4:**
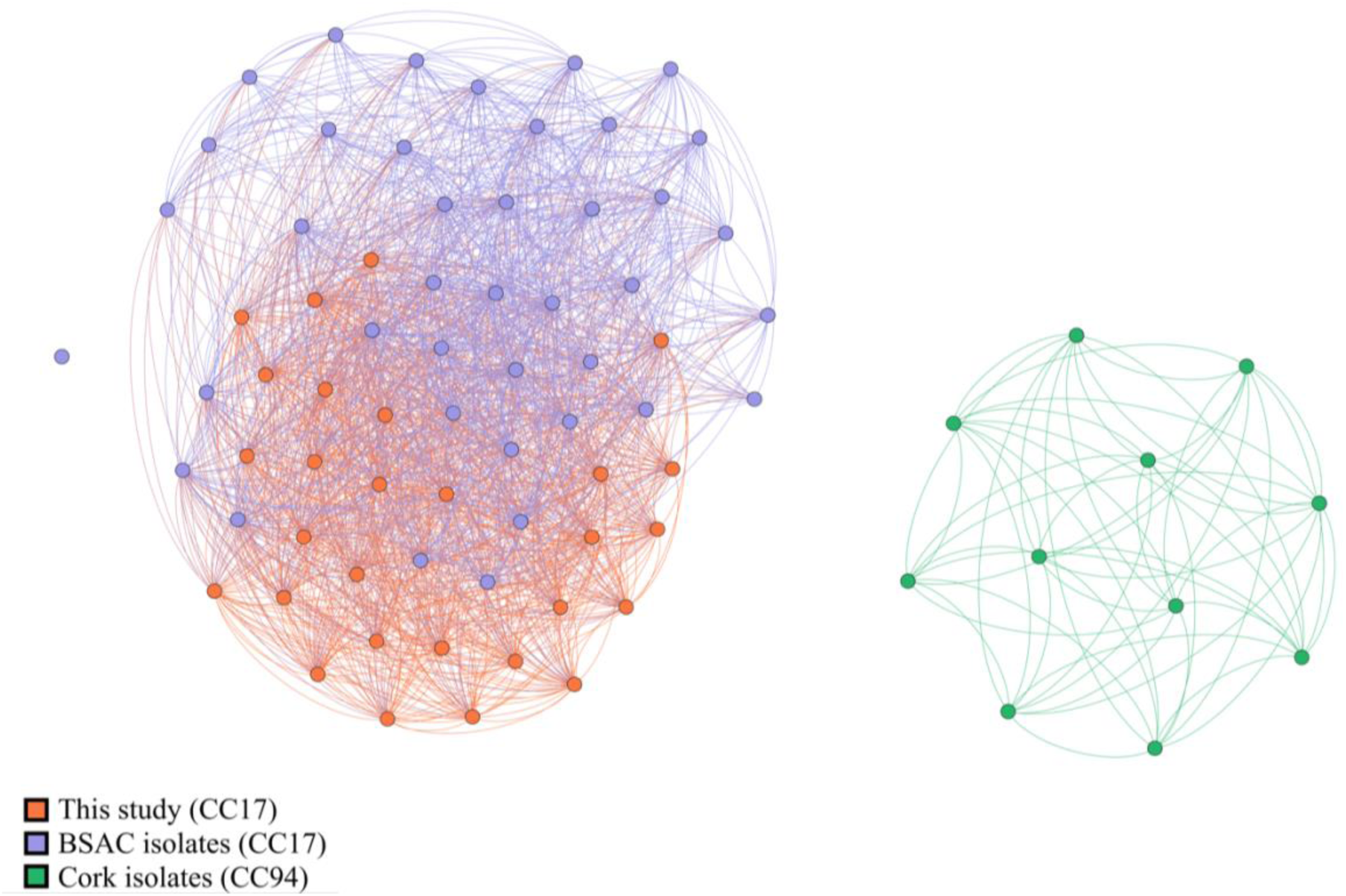
Genomic distance cluster analysis of mobilomal sequences from all isolates. While some mixing occurs with BSAC isolates and isolates sequenced for this study, divergence between communities is observed. One BSAC isolate (ERR374724) is completely disconnected from the other connected components, which is unusual due to the shared presence of a *vanA*+ genotype between ERR374724 and many other isolates. All CC94 isolates cluster together in a separate connected component from CC17 mobilomes.

### Pangenome analysis

A pangenome for the whole genome and chromosomal datasets (for all isolates, and individually for the “Cork” isolates, for the “BSAC” isolates, and for the isolates sequenced for this study) was produced using Roary *v.*3.1.3. (106) with the “-e” flag to align all gene clusters using PRANK *v.*170427 (107), and the “-z” flag to keep intermediate files (retained to produce the robust phylogeny below). To assess the effect of the mobilome on the genome diversity and complexity, each pangenomic category (*n*_genes_; “core pangenome” (99% ≤ *n*_samples_ ≤ 100%), “soft core pangenome” (95% ≤ *n*_samples_ < 99%), “shell pangenome” (15% ≤ *n*_samples_ < 95%), and “cloud pangenome” (*n*_samples_ < 15%)) in the whole genome generated pangenome was compared to the chromosomal pangenome using a two-tailed Fisher’s exact test (H_0_:ρ=π;H_A_:ρ≠π; (108)) with a Bonferroni-Dunn correction (*n*_comparisons_ = 4), instances where *P*_BD_ ≤ 0.05 were considered statistically significant, and statistically significant instances where ρ > π were considered to be overrepresented and instances where ρ < π were considered underrepresented (SI Table 13).

### Pangenome function

The representative pangenome refers to the collection of representative sequences from each pangenome cluster. Gene ontology terms (as assigned by InterProScan) were extracted and slimmed from the whole genome dataset (inclusive of all isolates) using the “map_to_slim.py” from GOATools (109) using generic .obo files. Sequences were grouped into their pangenomic categories and compared to the background population using the “find_enrichment.py” script (using a Fisher’s exact test (H_0_:ρ=π;H_A_:ρ≠π) and Bonferroni-Dunn correction (SI Table 14))

### Phylogeny construction

A phylogeny was constructed using single-copy, ubiquitous gene alignments (as produced by PRANK during pangenome construction) from the chromosomal dataset. The chromosomal dataset was used to further minimise the likelihood of interference non-vertically inherited sequences on the phylogeny. Each alignment was quality trimmed using TrimAL *v.*1.4 (110) using the “-automated1” flag. A superalignment was constructed by concatenating all trimmed alignments using FASconCAT *v.*1.04 (111) and a consensus phylogeny was constructed using IQtree *v.*1.6.12 (112) with 10,000 bootstrap replicates. IQTree used

ModelTest-NG *v.*0.1.7 (113) to determine the GTR+F+R6 (114) nucleotide evolutionary model to be most appropriate for phylogenetic reconstruction. The root of the tree was determined by creating a second phylogeny with a specified outgroup of 2 *Staphylococcus aureus* genomes. The second phylogeny was constructed by clustering all chromosomal and staphylococcal proteins using ProteinOrtho *v*.6.0.24 (115) with a stringency cut off value of *E*≤1.00*e*^-50^. All protein clusters that were ubiquitous in all *Enterococcus faecium* species and single copy for each species within each cluster were extracted and aligned using Muscle *v*.3.8.1551 (116). Each alignment was trimmed as above using TrimAL (using the “-automated1” flag) and a consensus tree was constructed as above using IQtree with 10,000 bootstrap replicates. ModelFinder-NG determined LG+I+G (117) to be the best model of protein evolution. The second phylogeny determined ST178 (all samples isolated in Cork, Ireland) to be the outgroup of the core gene tree. The finalised phylogeny was displayed and annotated using iToL *v.*5 (118) (Figure 5). The phylogeny was annotated with genome annotation data (generated using the analyses above and below) to visualise underlying trends (Figures 6-10; SI Figure 1)

**Figure 5:**
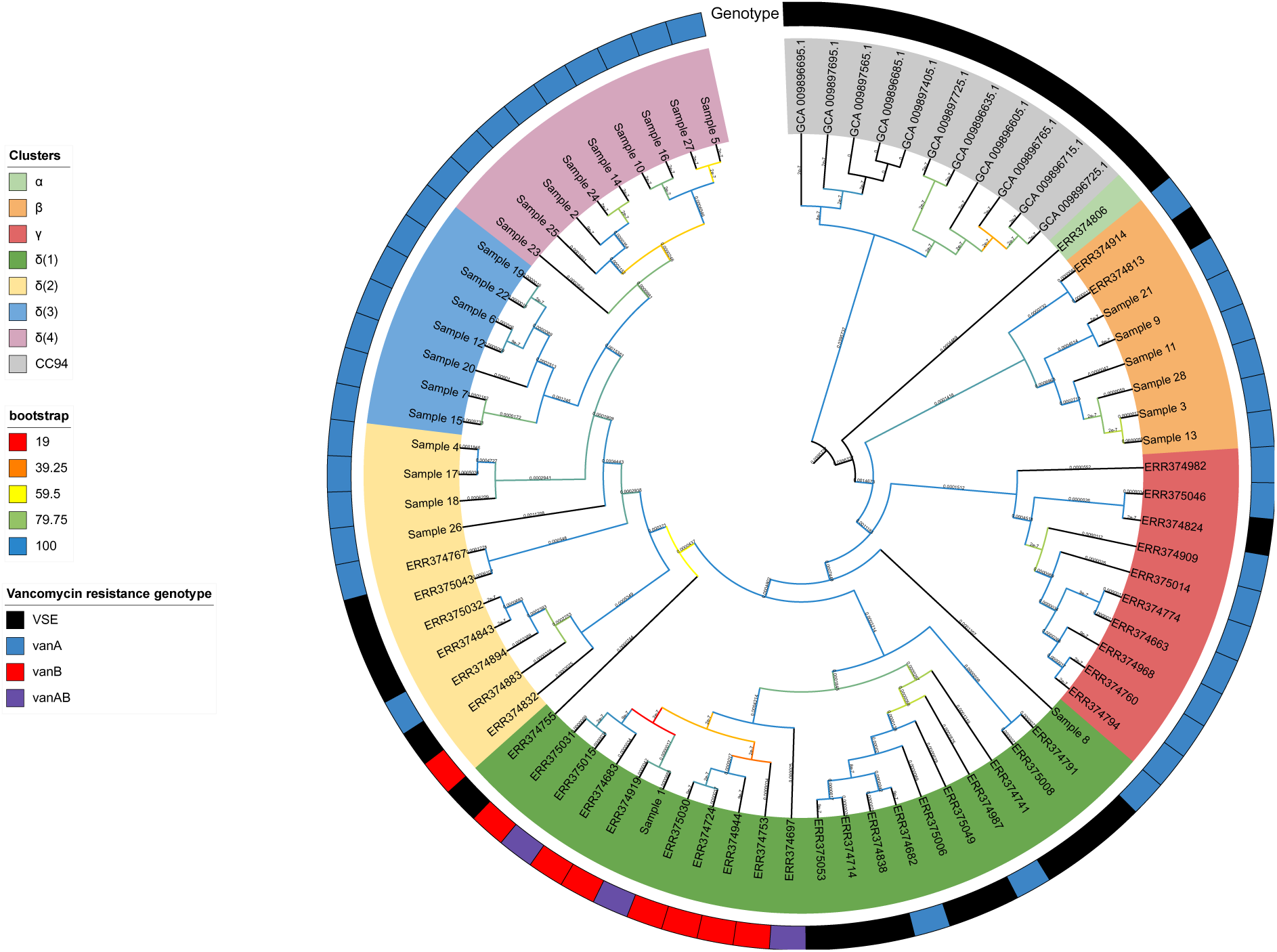
A 10,000 bootstrap phylogeny of isolates. Taxa are shaded based on their RheirBAPS clustering (discussed in methodology section) with an outer ring denoting the associated vancomycin resistance genotypes for each taxon. Bootstrap supports are provided for internal nodes. The majority of supports equate to 100%. Bootstrap shading is transitional, the colours shown in the legend are equidistant scales denoting where a new colour is shown. For example, a datapoint between 79.25% and 100% would be a shade of turquoise. Deep nodes all received high bootstrap support, further corroborating RheirBAPS clustering.

**Figure 6.**
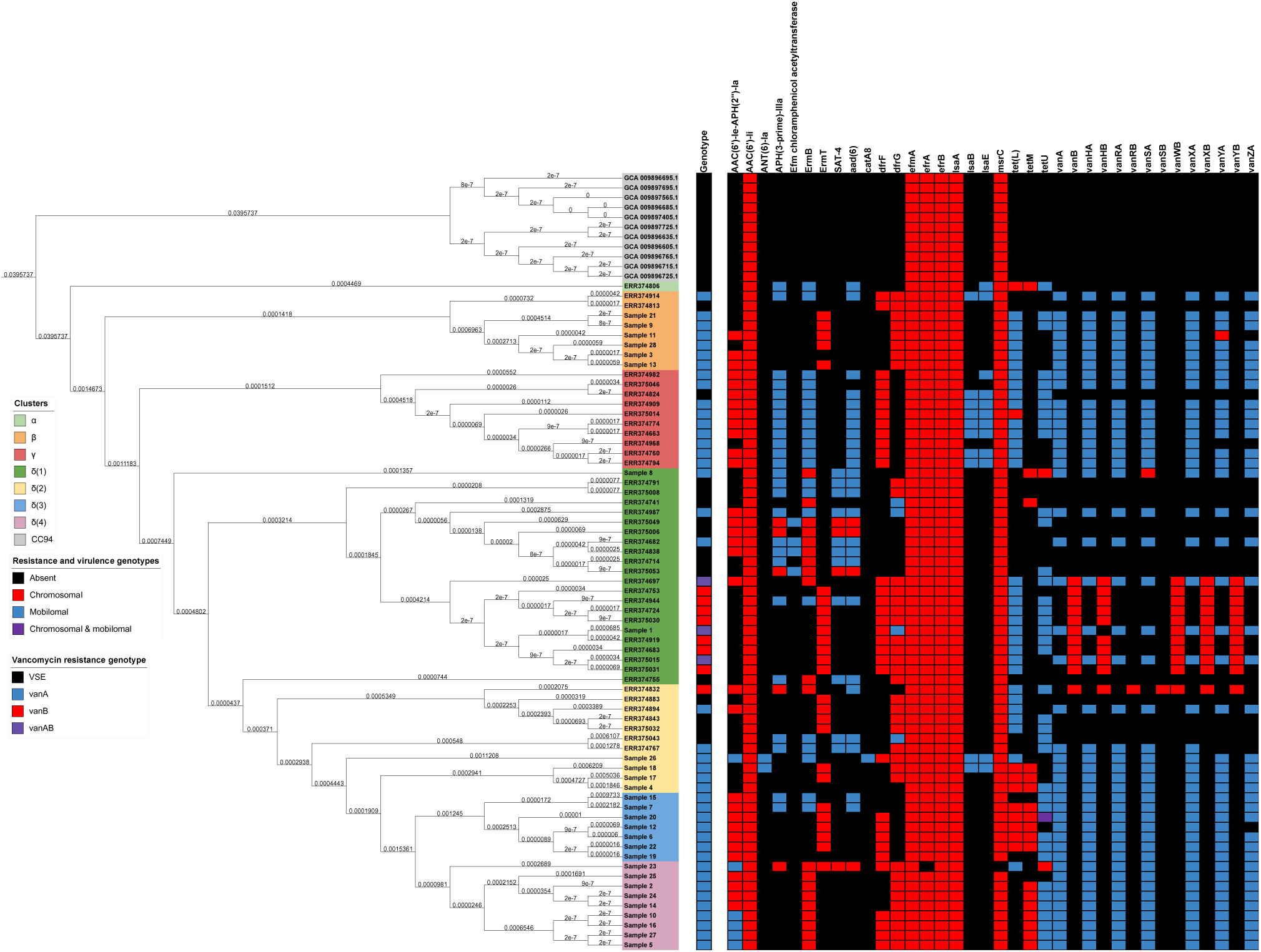
A phylogeny of isolates illustrating drug resistance genotype distributions as a categorical heatmap. Again, taxa are shaded based on their associated RheirBAPS cluster and a vancomycin resistance genotype is given as a separate bar for each taxon.

**Figure 7.**
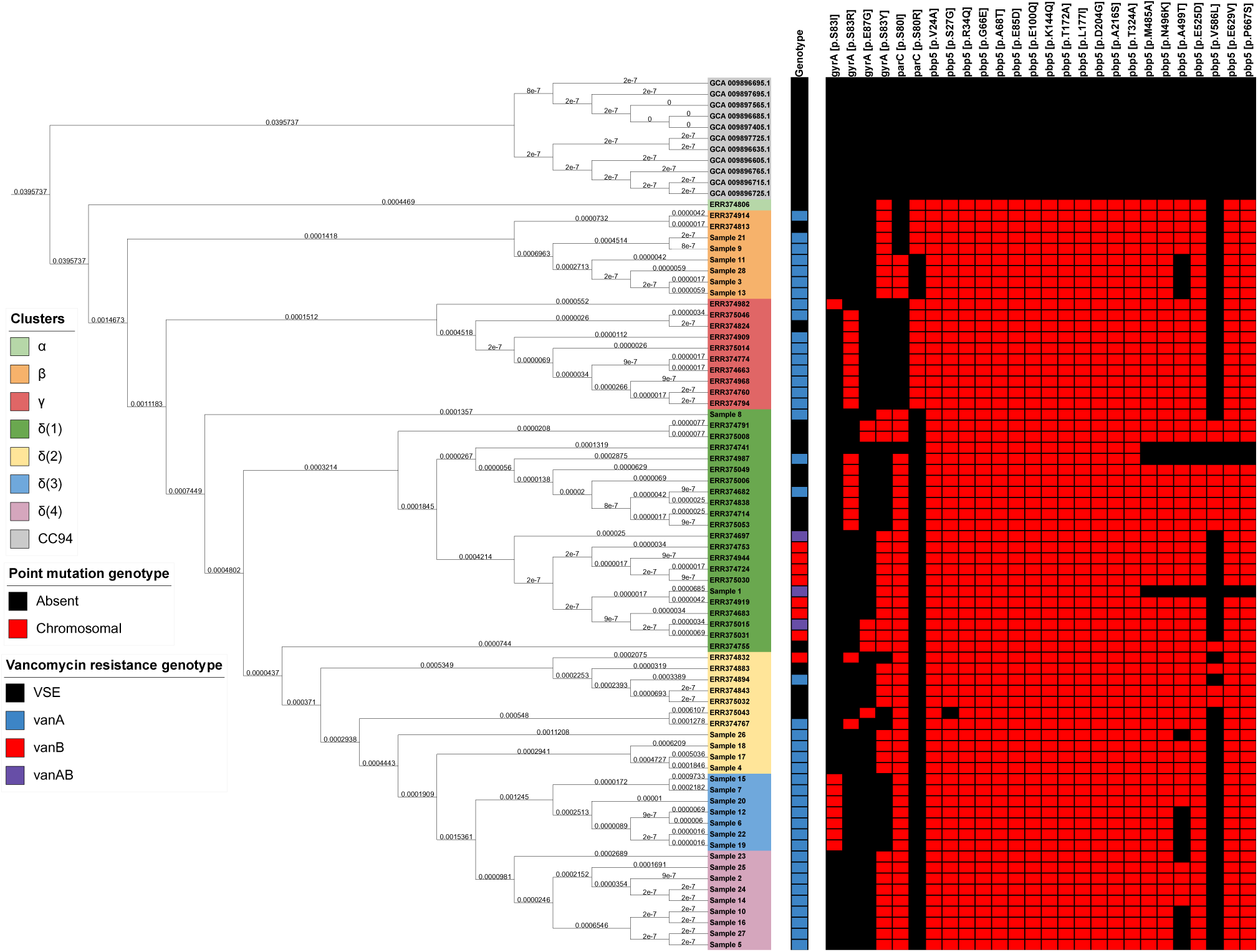
A phylogeny of isolates illustrating point mutation mediated drug resistance genotype distributions as a categorical heatmap. Again, taxa are shaded based on their associated RheirBAPS cluster and a vancomycin resistance genotype is given as a separate bar for each taxon.

**Figure 8.**
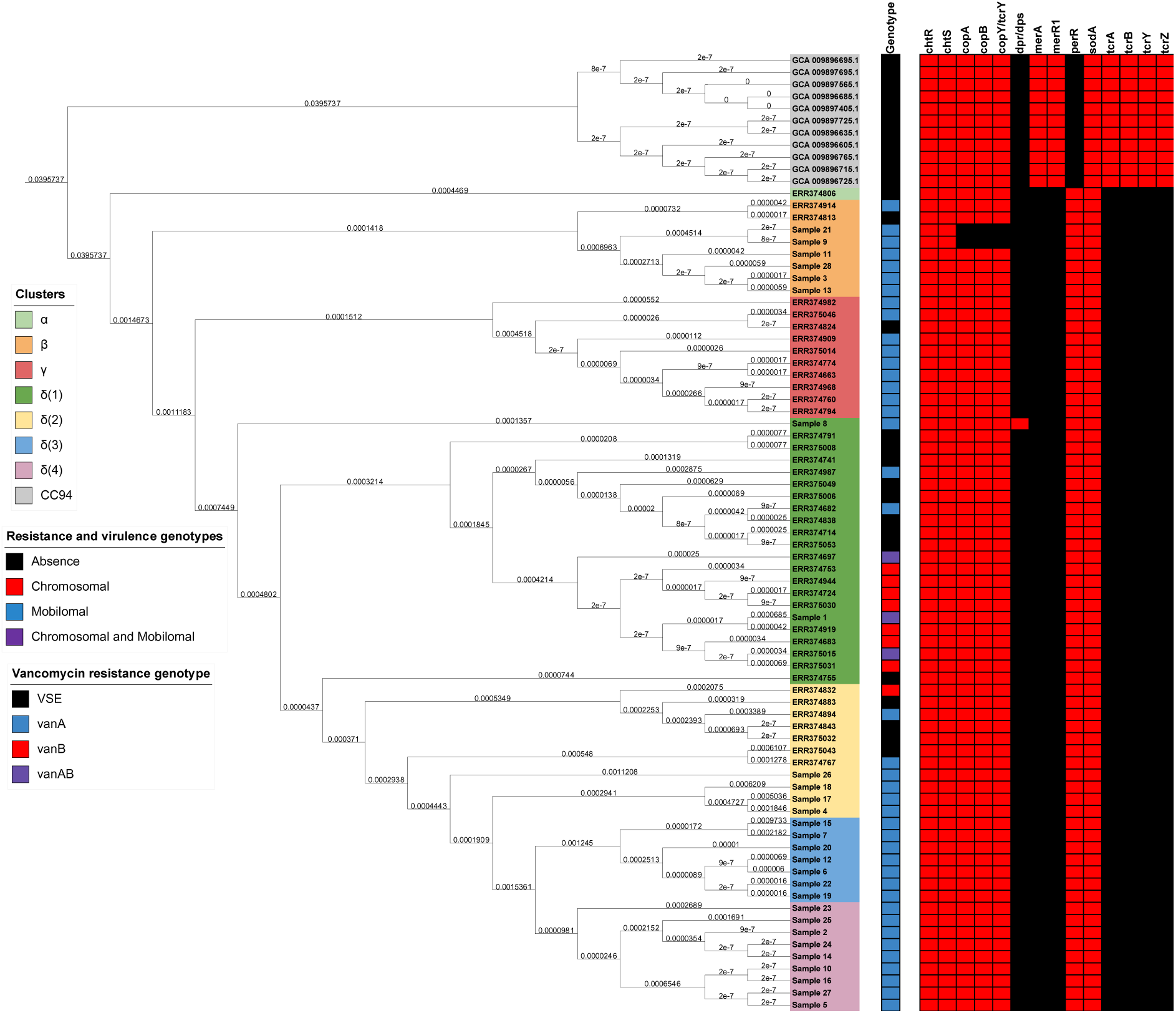
A phylogeny of isolates illustrating metal (biocide) resistance genotype distributions as a categorical heatmap. Again, taxa are shaded based on their associated RheirBAPS cluster and a vancomycin resistance genotype is given as a separate bar for each taxon.

**Figure 9.**
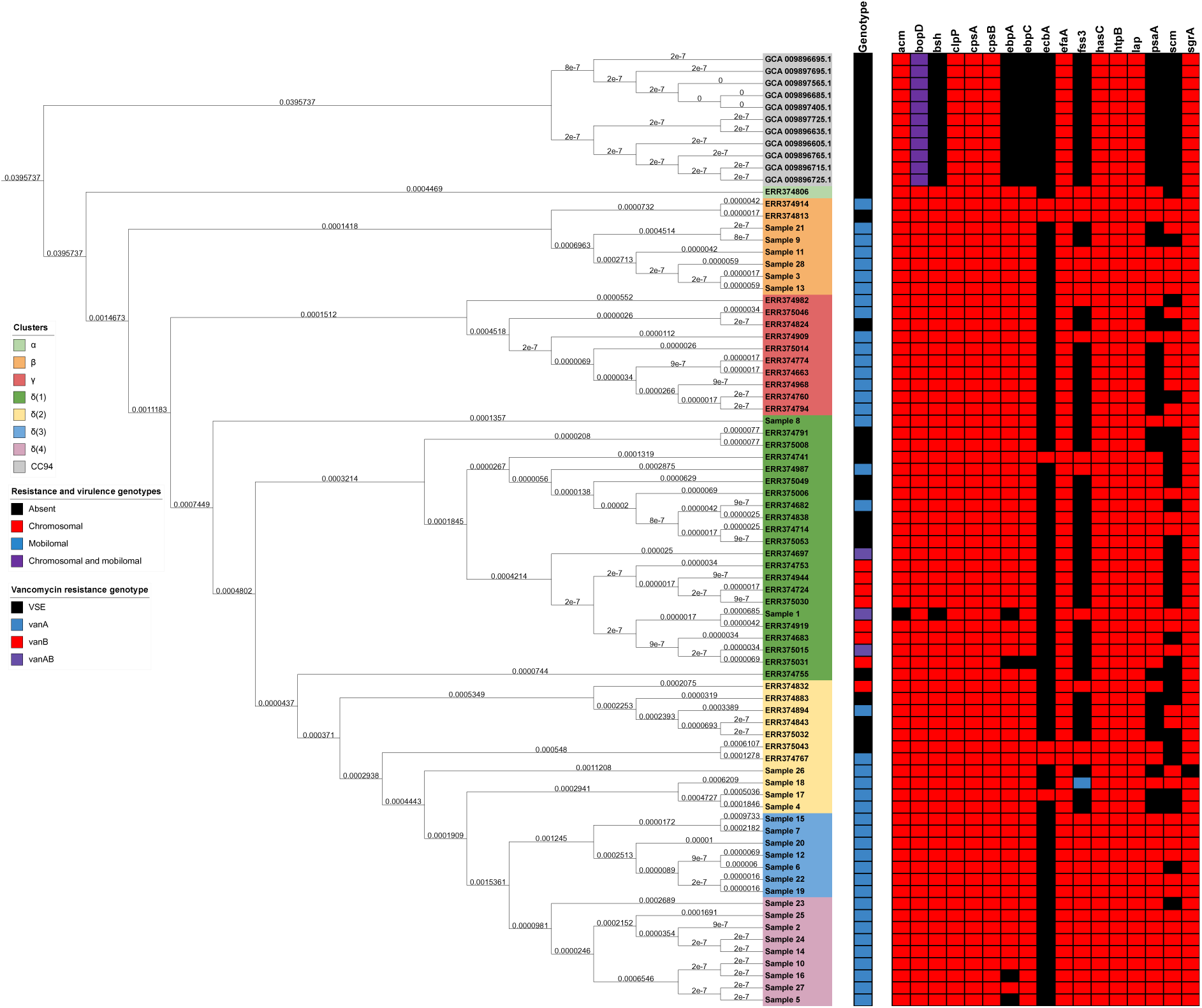
A phylogeny of isolates illustrating virulence factor genotype distributions as a categorical heatmap. Again, taxa are shaded based on their associated RheirBAPS cluster and a vancomycin resistance genotype is given as a separate bar for each taxon.

**Figure 10.**
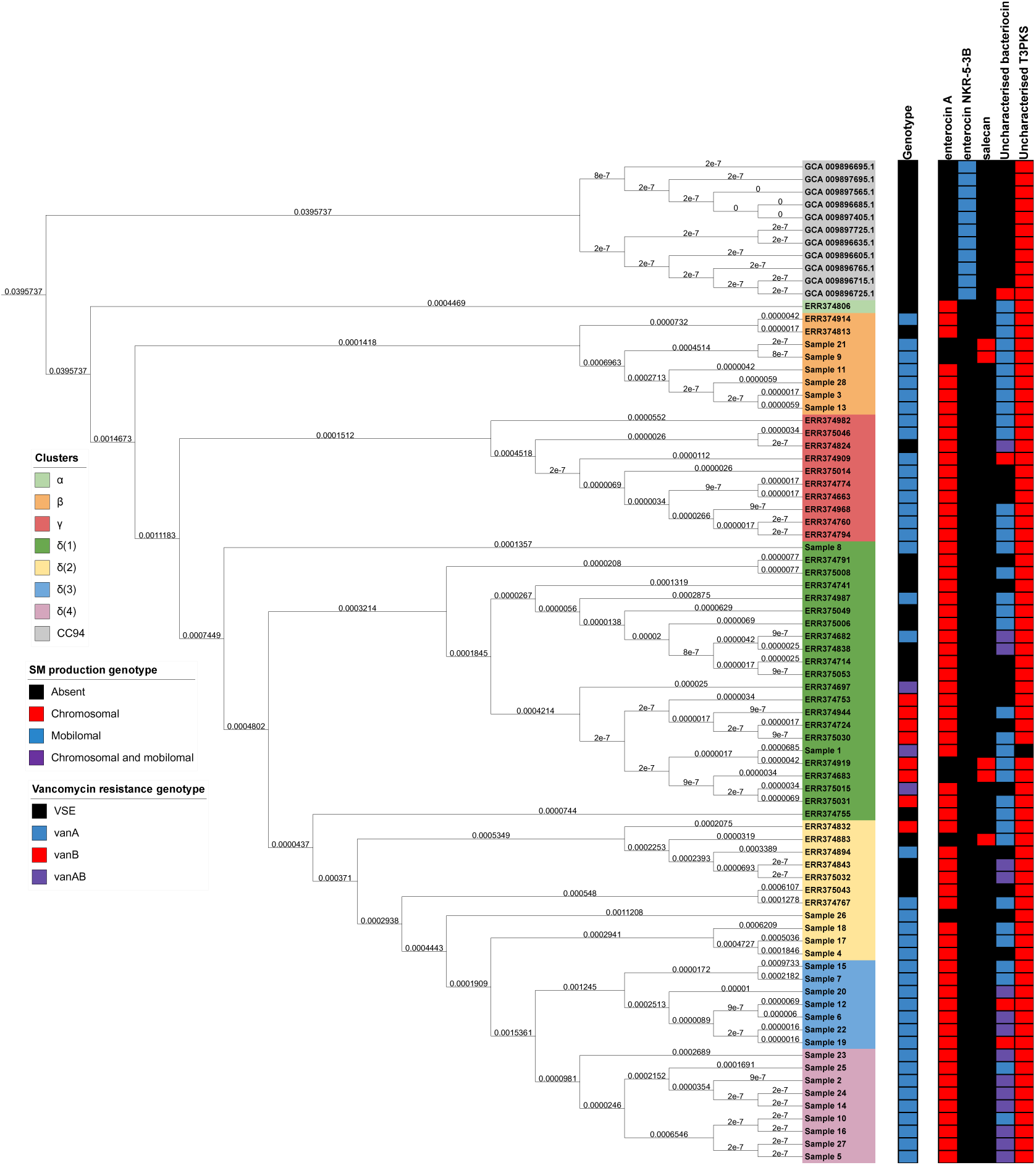
A phylogeny of isolates illustrating secondary metabolite production genotype distributions as a categorical heatmap. Again, taxa are shaded based on their associated RheirBAPS cluster and a vancomycin resistance genotype is given as a separate bar for each taxon.

### Isolate clustering

Isolates were clustered using previously constructed core gene superalignment using RheirBAPS *v.*1.1.3 (119, 120) with a maximum depth of two, an initial population cluster of 20, and with an additional 100 rounds of processing to approximate optimal clustering (Figure 5; Figure 11; SI Table 15). All isolate-cluster assignments were determined to approach 100% likelihood (*P* = 1 in all cases) indicating that their placement is correct.

**Figure 11:**
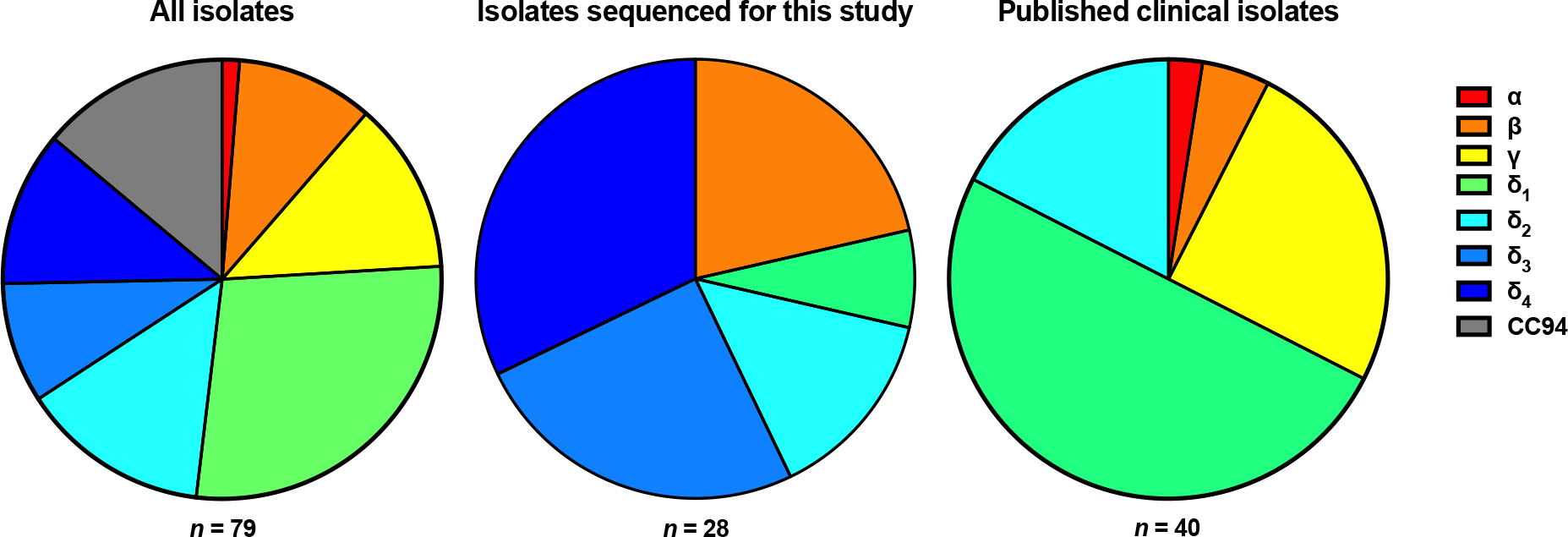
Distribution of RheirBAPS clusters between studies

### Core genome MLST

A core genome MLST (cgMLST) was constructed for all isolates using chewBBACA *v.*2.8.5 (121) using default settings and using the *Enterococcus faecium* Prodigal training file provided with the software. The cgMLST profile was displayed and annotated with metadata using GrapeTree (122) with the MSTree V2 algorithm (Figure 12). The generated cgMLST was verified using our previously generated data pangenomic data with PANINI *v*.1 (123). Briefly, as PANINI requires a minimum of 100 input taxa, we constructed a pseudo-dataset where each taxon was represented twice and processed. Pseudo-taxa were removed from the PANINI output and was visualised alongside the previously constructed phylogeny and annotated with their associated RheirBAPS generated clade using MicroReact (124) (SI Figure 2).

**Figure 12:**
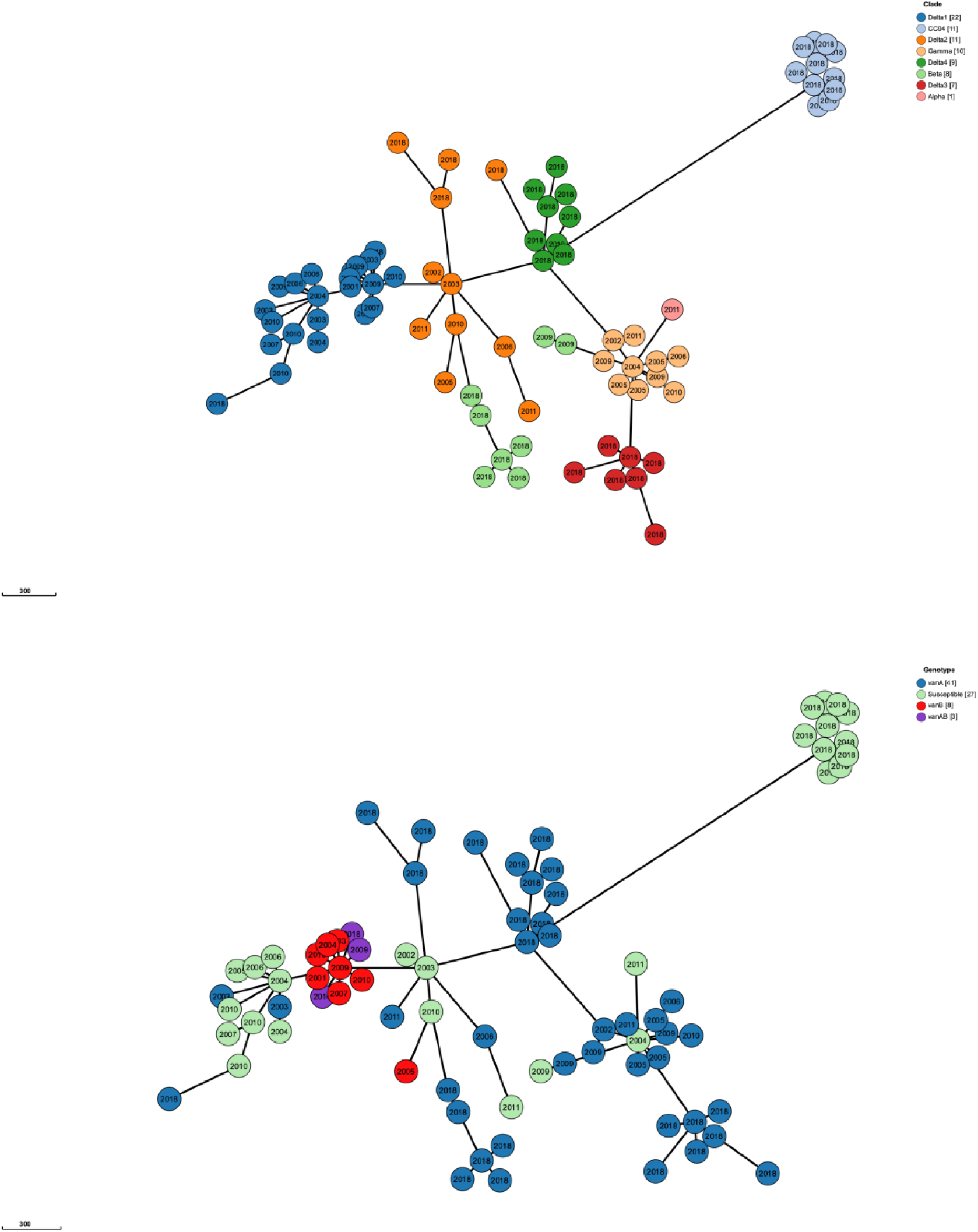
Core genome MLST (cgMLST) for all isolates included in this study. The upper image is shaded based on RheirBAPS clustering and the lower is shaded based on vancomycin resistance genotypes. The year of isolation is presented in the centre of each circle, where each circle represents an isolate.

### Pangenomic co-evolutionary analyses

The chromosomal phylogeny and Roary-derived pangenome (inclusive of all isolates) were used to test statistically significant associations or dissociations between any vancomycin resistance gene (*vanA, vanB, vanH, vanW, vanX, vanXB, vanY, vanYB*) and any other gene (SI Table 16) using Coinfinder (56). Coinfinder uses a binomial test (H_0_:π=π*_x_*;H_A_:π≠π*_x_*) and while the authors advise the use of a Bonferroni-Dunn corrected, this was not implemented so all coincidences could be explored and instances where *P* ≤ 0.005 were considered statistically significant.

### Recombination analysis

Using RheirBAPS resulted into two distinct clades, the “Cork” isolates (CC94) and all other isolates (CC17; discussed in a later section). To examine the extent of core genome recombination on CC17 evolution, genomic comparisons were performed between each CC17 isolate and a “complete genome” *Enterococcus faecium* (reference) assembly (downloaded from NCBI Assembly). The reference was selected from a pool of all available *Enterococcus faecium* complete genome assemblies (which passed taxonomy assignment) from NCBI Assembly, where each genome was assessed for *vanA* or *vanB* presence using ABRicate with the CARD database (--minid 50), assigned a sequence type using MLST, and assigned a CC using PubMLST. Any assembly with a *vanA*+ or *vanB*+ phenotype, with an indeterminant ST, or assigned to either CC17 or CC94 were discarded. Remaining isolates were processed to remove plasmids and the remaining chromosomal sequence was annotated using Prokka with default settings. Proteins from all CC17, CC94, and candidate reference assemblies were clustered using ProteinOrtho (*E*≤1*e*^-50^). Protein clusters that were both single copy and ubiquitous in all species were individually aligned using Muscle with uninformative regions using TrimAL (with the “-automated1” flag) and concatenated into a superalignment using FASConCAT. The superalignment was processed to construct a consensus tree with 10,000 bootstrap replicates using IQTree. The resultant consensus tree was rooted at CC94 and the closest assembly to CC17 was selected as the reference strain. The most suitable reference was determined to be GCF_005166365.1 (strain NM213; PRJNA513159). Strain NM213 was isolated from healthy Egyptian infants in 2018 to determine its potential as a probiotic (https://www.ncbi.nlm.nih.gov/bioproject/PRJNA513159/).

Whole genome alignments were constructed between strain NM213 and each CC17 assembly and concatenated using Snippy *v.*4.6.0 (snippy-multi) with default settings (https://github.com/tseemann/snippy). The resultant core whole multigenome alignment was processed to convert non-standard nucleotide characters to (ATGC) to “N” using a Snippy auxillary script (“snippy-clean_full_aln”). Genomic recombination regions were detected using Gubbins *v.*3.0.0 (125) using default settings and the extent of recombination was illustrated using Phandango *v.*1.3.0 (126) where the previously constructed core-genome phylogeny (of CC17 and CC94) was used to scaffold recombination (Figure 13). This procedure was replicated for the remaining 24 suitable references (SI Figure 3).

**Figure 13:**
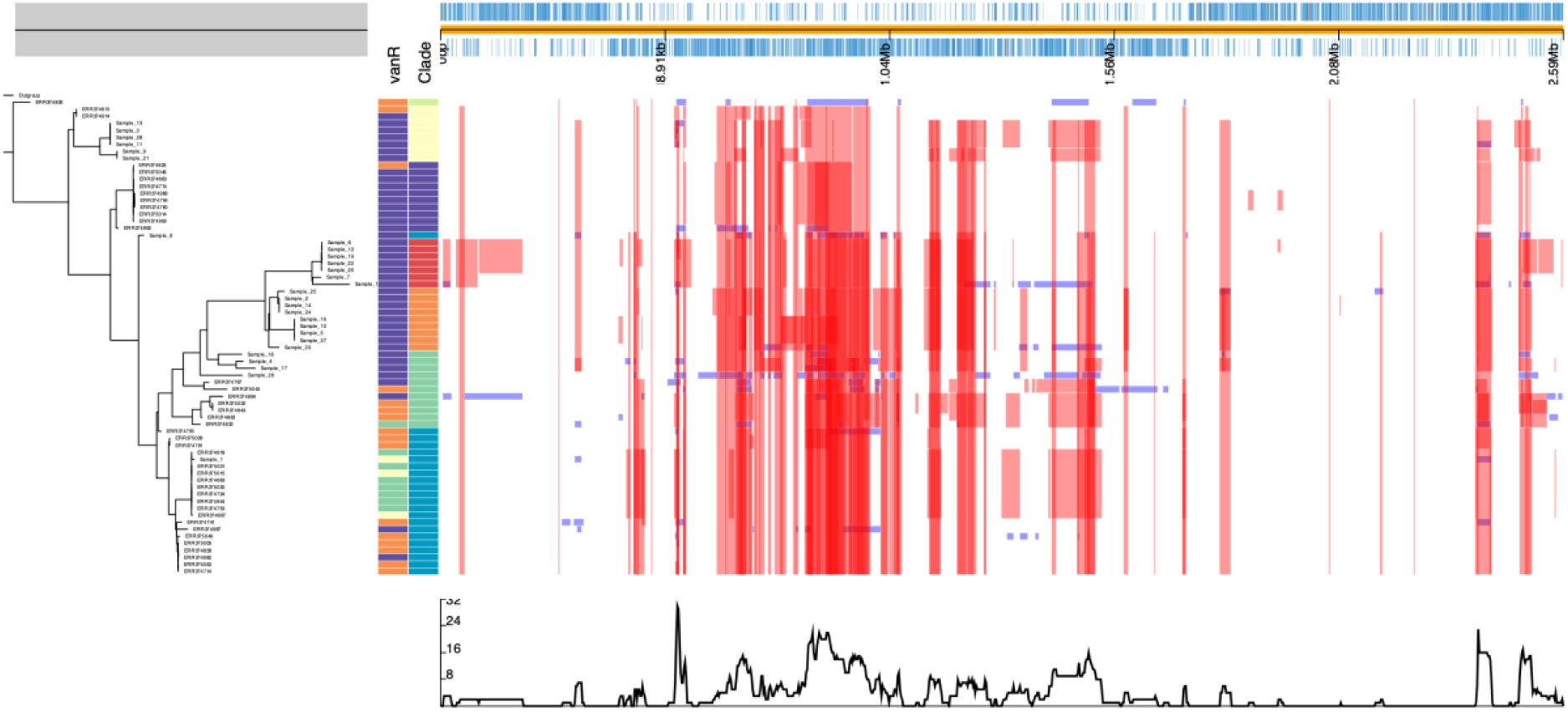
Red bars illustrate hot spots compared to a reference, where a hotspot is defined as a site where more than one species reports a ≥50% likelihood for the same recombination event. Blue boxes illustrate a recombination event for a single species. Multiple different recombination events can occur at the same site (appearing as overlap in this image). The plot at the bottom shows the maximal amount of species affected by a detected recombination event in a given hotspot

### Phages

Phage and prophage elements within each chromosomal and plasmid sequence were assessed using Phigaro *v.* 2.3.0 (127) with default settings and saving the detected viral sequences to their own output files (“--save-fasta” flag) (SI Table 17). The sum of phages per genome were counted (SI Table 18). Extracted viral sequences were assessed as a source of drug resistance, metal resistance and virulence factors using abricate with the CARD, BACMET, VFDB databases as above and as a source of secondary metabolism machinery using GECCO as above, only one gene-of-interest was observed (discussed below). Extracted prophages were annotated using Prokka using the “--kingdom Viruses” flag.

### Phage classification

A database of all Caudoviridae ICTV exemplar reference phage genomes were downloaded from NCBI Assembly. Caudoviridae were selected as this clade encompassed all viral families identified by Phigaro (Siphoviridae, Inoviridae, and Myoviridae, respectively). Each extracted prophage was searched against the database using Mash *v.*2.2.2 (75) using the “dist” (distance) algorithm. As we searched prophages (partial genomes) against phages (whole genomes) we selected top hits (1 per prophage) from instances with the shortest distances (*D* < 1) and smallest *P*-values (*P* ≤ 0.005) for each prophage (SI Table 18; Figure 14).

**Figure 14.**
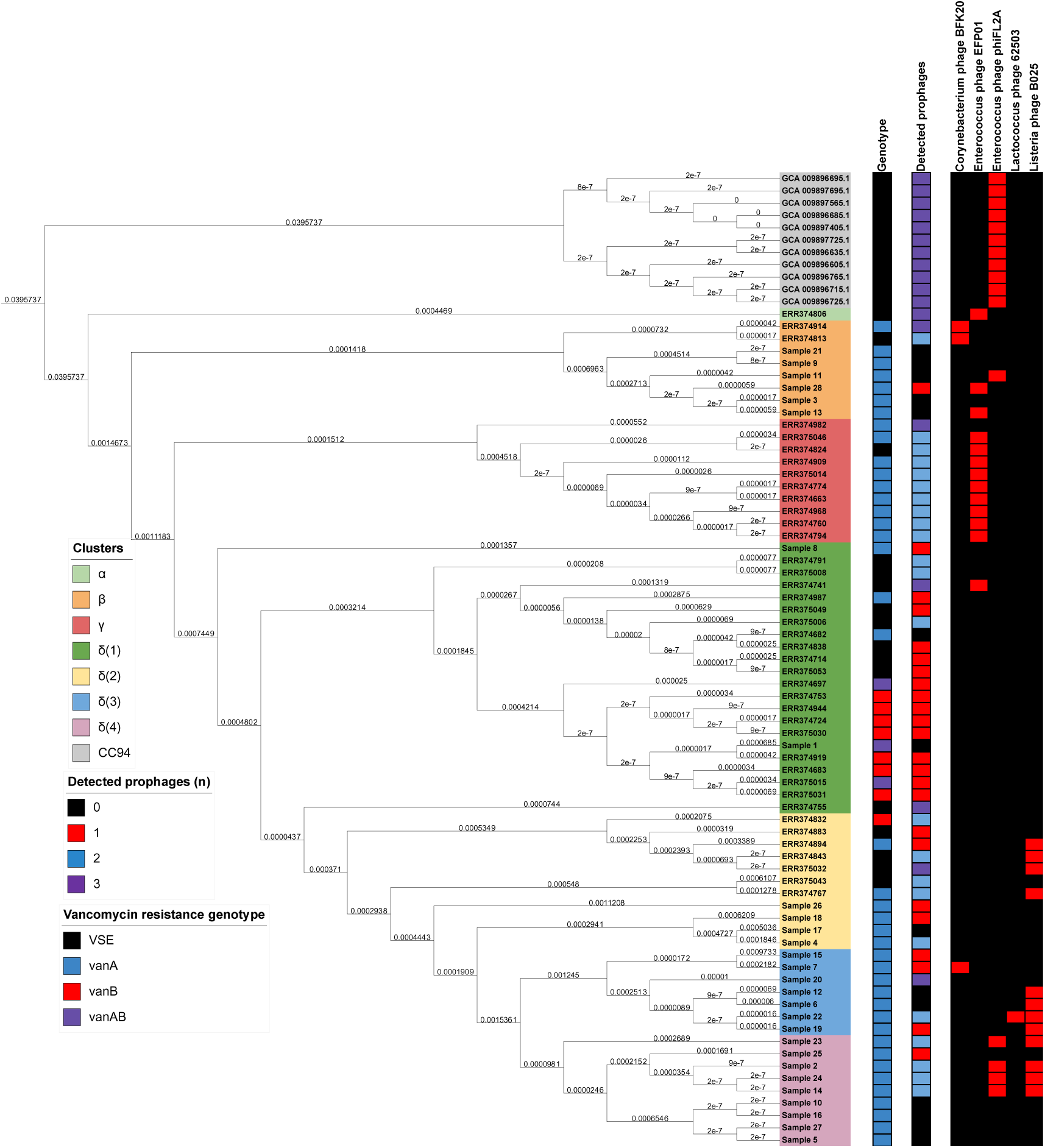
A phylogeny of isolates illustrating detected prophage distributions as a categorical heatmap. Again, taxa are shaded based on their associated RheirBAPS cluster and a vancomycin resistance genotype is given as a separate bar for each taxon.

### Integron assessment

The presence of integrons in chromosomal and plasmid sequences was determined using IntegronFinder *v.*2.0 (128) using the “--local-max” flag and setting “--calin-threshold” flag to 1 (instead of the default 2). We chose to set the “--calin-threshold” flag to 1 to include integrated regions with CALIN artefacts. While lowering this threshold may yield a higher number of false positives, we feel it is appropriate for exploratory purposes. The distribution of integrons per genome is given in SI Table 19. Again, ABRicate was used to determine whether contigs containing integron elements contained genes-of-interest (SI Table 20; Figure 15).

**Figure 15.**
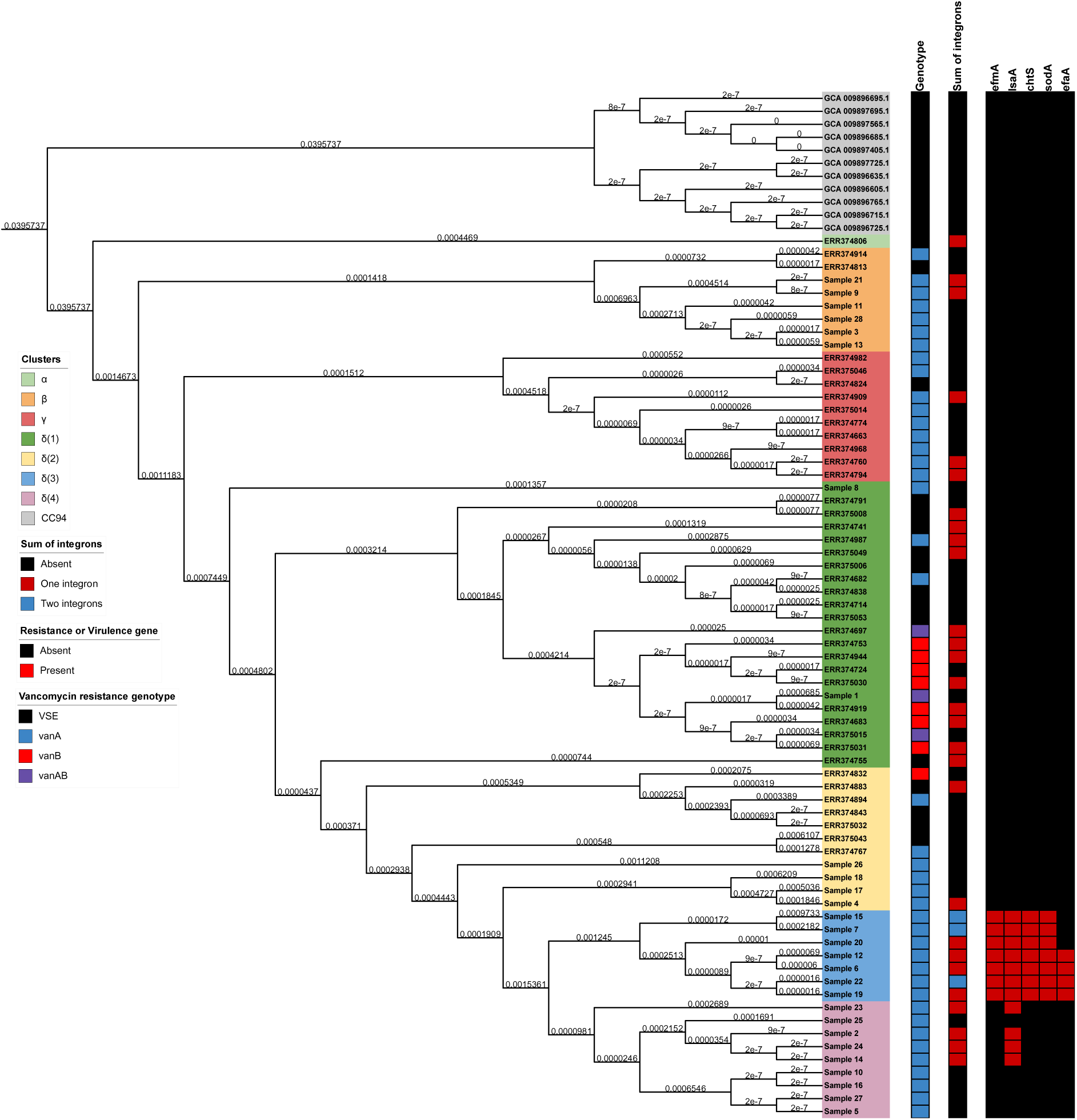
A phylogeny of isolates illustrating detected integron distributions as a categorical heatmap. Again, taxa are shaded based on their associated RheirBAPS cluster and a vancomycin resistance genotype is given as a separate bar for each taxon. Gene distributions (presence or absence) are given for *efmA* and *lsaA* (drug resistance), *chtS* and *sodA* (metal and biocide resistance), and *efaA* (virulence factor).

## Results

### Quality control

All assemblies were reported to have a high level of completeness (μ_completeness_ = 99.29±0.55%; 95.16 ≤ % ≤ 99.61; SI Table 3).

### General characteristics of the genomes

Genomes used in this manuscript were reported to have a mean chromosomal gene count of 2537.91±77.44 (η = 2550.5; 2334 ≤ *n* ≤ 2730) and an aggregated (chromosomal and mobilomal) gene count of 2,821.64±116.65 (η = 2812; 2508 ≤ *n* ≤ 3158). For chromosomal data, a mean gene sum of 2537.91±77.44, a mean genome size of 2681.5±78.77 Mbp, a mean density of 0.946±0.0098 genes/Mbp, and a mean GC% of 38.03±0.15% (SI Tables 10-11; Figure 2). For mobilomal data, a mean gene sum of 283.73±84.86, a mean genome size of 259.99±75.76 Mbp, a mean density of 1.088 genes/Mbp, and a mean GC% of 34.70±0.38%. When statistically examined, mobilomes were observed to be significantly denser with regards the number of observed genes (to have a greater number of genes per Mbp genome; *P*_BD_ < 1.00*e*^-06^) and had a significantly lower GC% (*P*_BD_ < 1.00*e*^-06^) When the coding proportions were compared, 19 mobilome samples were reported to be significantly less dense (in terms of the sum of coding nucleotides) than their chromosomal counterparts, however 61 comparisons displayed insignificant differences.

### Genetic relatedness (MLST, cgMLST, and RheirBAPS)

While a total of 16 STs were observed throughout the dataset, considerable differences were observed between the 3 sources (isolates sequenced for this study, previously published isolates, and “Cork” isolates). The “Cork” isolates were exclusively ST178, and ST178 was not observed in either of the clinical sources (Figure 1, SI Table 4). Isolates sequenced for this study were predominantly ST80 (16 of 28; 57.14%) and previously published isolates were predominantly ST17 (17 of 40; 42.5%) and ST192 (9 of 40; 22.5%). Interestingly, ST80 and ST17 are exclusively observed in their associated sources, however ST192 and ST203 are observed in both sources.

Isolate clustering using RheirBAPS matched clonal complexes at level one, with all CC94 isolates assigned to Cluster A and all CC17 isolates assigned to Cluster B (SI Table 14). At level two, eight clusters were observed, however using phylogenetic inference, this can be collapsed to five clusters with four subclusters. As all CC94 isolates were again assigned to a single cluster, CC17 isolates formed four clusters (α, β, γ, δ) with four subclusters (δ_1_, δ_2_, δ_3_, and δ_4_). These clusters are named based on their phylogenetic divergences (Figure 5) . Cluster α contained one genome, ERR374806 (ST1032) and was the earliest diverging CC17 isolate in our dataset. Cluster β encompassed all ST203 isolates and all ST192 isolates except for ERR374982 (which was assigned to Cluster γ, matching its phylogenetic placement). Cluster γ encompasses all ST78 isolates and the singleton ST192. As previously mentioned, Cluster δ was comprised of four distinct subclusters (δ_1_, δ_2_, δ_3_, and δ_4_). Where δ_1_ is basal to all other δ clades, and δ_2_ is basal to sister clades δ_3_ and δ_4_. Clade δ_1_ encompassed all ST17 isolates, and isolates assigned to singleton ST groups (ST18, ST132, ST982, ST1038, ST1421). Clade δ_2_ encompassed all ST80 isolates, two ST80 isolates, and singleton ST isolates (ST16, ST64, ST132, ST202, ST612). Clade δ_3_ encompasses seven ST80 isolates and clade δ_4_ encompasses the remaining eight ST80 isolates and the singleton ST787. These results suggest that ST80 is a descendant of ST17 and is undergoing considerable chromosomal evolution. Clusters α, β, and γ, and Subclusters δ_3_, and δ_4_ all displayed monophyly. Subcluster δ_2_ displayed considerable paraphyly with 4 root nodes. Subcluster δ_1_ also displayed paraphyly (if the Cluster δ earliest diverging taxon (Sample 8) is ignored) due to ERR374755. The Subcluster δ_1_ paraphyly may be considered an artefact of cluster evolution, with Subclusters δ_2_, δ_3_, and δ_4_ undergoing rapid evolution after the divergence of ERR374755 from their common ancestor. The paraphyly observed in Subcluster δ_2_ may be due to sample size and may be resolved or further partitioned with additional taxa. The paraphyly observed is likely due to evolutionary expansion between sampling times, with earlier diverging isolates (BSAC isolates) being isolated prior to 2011 and later diverging isolates (isolates sequenced for this study) being sequenced in 2018).

The cgMLST graph (and associated graph analyses) displayed considerable modularity (Figure 12; SI Figure 2) with distinct clades forming, largely corroborating RheirBAPS results, regardless of the year of isolation. Two exceptions were observed for this trend: firstly, polyphyly in Cluster β was observed with isolates from 2009 adjacent to Cluster γ, and those from 2018 adjacent to Subcluster δ_2._ Again, Subcluster δ_2_ appeared less modular than other Clusters (mirroring paraphyly observed in the phylogeny), but this was likely due to core genome evolution between sampling dates. A Subcluster δ_2_ isolate was observed adjacent to Subcluster δ_4_.

### Plasmid containment

As expected, the majority of PLSDB hits were attributed to *E. faecium* (SI Table 5), however some other interesting observations were also noted. All eleven CC94 (Cork) isolates each displayed a hit to two *Listeria monocytogenes* strain CFSAN023459 plasmids (pCFSAN023459_01 and pCFSAN023459_02), eight CC94 isolates also returned a hit for *Enterococcus hirae* strain CQP3-9 plasmid pCQP3-9_1. Hits pertaining to *Bacillus cereus* plasmid pBC16 were observed across CC17, interestingly, all Cluster γ isolates except ERR375014 (ST78) and all Cluster β isolates except ERR374813 and ERR374914 (ST203; only Cluster β isolates not sequenced during this study) returned a hit. Plasmid pEGM182-2 from *Enterococcus casseliflavus* strain EGM182 was observed in CC94 and in Subclusters δ_3_ and δ_4_, where four of six δ_3_ isolates (except Samples 6 and 12; ST80) returned two hits for pEGM182-2. Hits were returned for *Escherichia coli* strain UB-ESBL31 plasmid pESBL31 from isolates across the phylogeny. Of note, all Cluster γ isolates and five δ_4_ isolates returned a hit for pESBL31. Most of these isolates (14 of 15) were sampled between 2002 and 2011, and one isolate (Sample 1) was sampled in 2018, suggesting persistent survivability for this plasmid and its descendants.

### Antimicrobial susceptibility profiles

The 28 vancomycin resistant *Enterococcus faecium* were isolated from 22 hospital patients in 2018/2019 (MMUH, Dublin isolates). Samples 2 and 24 were isolated on the same day from patient 2. Samples 10, 5 and 27 were isolated from patient 5 at days 92, 100 and 107, respectively. Samples 21, 20, 9 and 26 were isolated from patient 9 on days 0, 8, 19 and 365, respectively. The antimicrobial susceptibility profiles (SI Table 2) of the isolates confirmed they were all vancomycin resistant. Of the 28 VRE*fm*, 25 were erythromycin resistant, two were chloramphenicol resistant, 28 were ampicillin resistant, 19 were tetracycline resistant and two were linezolid resistant. Samples 14 and 24 were the only isolates resistant to chloramphenicol or linezolid. While sample 2 was isolated from the same patient as sample 24 it did not display the same resistance profile. The three samples susceptible to erythromycin did not display a specific AMR gene absence associated only with these samples. The metadata of the BSAC study contained susceptibility to vancomycin data only and the “Cork” isolates contained no antimicrobial resistance data. Of the BSAC study isolates 23 were vancomycin susceptible and 17 were vancomycin resistant (SI Table 1).

### Antimicrobial resistance genotyping

A total of 32 resistance genes were observed throughout the dataset, of which 13 were exclusively observed in the chromosome (AAC(6’)-Ii, *dfrF, ermT, efmA, msrC, tetM, vanB, vanHB, vanRB, vanSB, vanWB, vanXB,* and *vanYB*), where one pair (*vanRB* and *vanSB*) were observed exclusively in isolate ERR374834. Ten genes were observed exclusively on the mobilome (ANT(6)-Ia, *Enterococcus faecium* chloramphenicol acetyltransferase, *catA8, lsaE, tetU, vanA, vanHA, vanRA, vanXA,* and *vanZA*), and 9 were observed in either the chromosome or mobilome (AAC(6’)-Ie-APH(2’’)-Ia, APH(3’)-IIIa, *ermB, SAT-4, aad*(6)*, dfrG, tet(L), vanSA*, and *vanYA*). Interestingly, the 9 genes present on either the chromosome or mobilome never appeared both chromosomally and mobilomally in the same isolate (SI Table 6; Figure 6).

### Mobile AMR genes

The *vanA* gene was present only in the mobilome of 44 isolates, almost always with the cluster of *vanHA*, *vanRA*, *vanSA*, *vanXA*, *vanYA* and *vanZA* (SI Table 6; Figure 6). These mobilomes were sampled from across CC17, spanning Clusters β, γ, and δ (encompassing 16 ST80 isolates, 7 ST78 isolates, 5 ST203 isolates, 4 ST17 isolates, and 3 ST192 genomes, and one isolate from each of ST16, ST18, ST64, ST132, ST202, ST612, ST1032, ST1421 and an unassigned ST).The *vanA* isolates comprised samples from the BSAC study (*n* = 16) isolated in 2002, 2003, 2005, 2006, 2007 and 2009 to 2011 and the isolates from Dublin between 2018 and 2019 (*n* = 28). The *vanB* gene was identified on the chromosomes in addition to the mobile *vanA* on two isolates from the BSAC study and one isolate from MMUH (sample 1). Samples 14 and 24 (ST80, Subcluster δ_4_) were linezolid resistant but did not contain the mobile linezolid resistance genes. Both samples were also the only isolates displaying chloramphenicol resistance. However, neither contained any known mobile chloramphenicol resistance genes. *Enterococcus faecium* chloramphenicol acetyltransferase (conferring resistance to phenicols) was observed in the mobilome of 3 of 17 ST17 isolates and in the ST1038 isolate (all Subcluster δ_4_). Another phenicol resistance gene *catA8* was also observed in the mobilome of sample 26. However, all were phenotypically susceptible to chloramphenicol.

The resistance gene *lsaE* (conferring resistance to lincosamide) was observed on the mobilomes of samples 18 and 26 and ten isolates from the BSAC study. The genes *aad*(*6*)*, ermB* and *aph(3’)III-a* were present in each of the BSAC mobilomes containing *lsaE*. The tetracycline resistance gene *tetU* was observed on the mobilome of 6 ST80 isolates, 2 ST18 isolates, and one isolate from each of ST17, ST132, ST787, ST1038, and ST1421 respectively. In addition, *tetL* (conferring resistance to tetracyclines) was frequently present on the mobilomes.

### Chromosomally mediated AMR

The *vanB* gene was identified in 11 isolates and were always chromosomal. A cluster of *vanB, vanWB, vanXB,* and *vanYB* (conferring resistance to glycopeptides, specifically vancomycin) was observed in nine Subcluster δ_1_ isolates (accounting for one ST16, and 8 of 17 ST17 isolates) and one Subcluster δ_2_ isolate (ERR374832; ST132) (SI Table 6; Figure 6). With the exception of ST16, all isolates with these four genes were also observed to possess *vanHB*, and the ST132 isolate was further observed to possess *vanRB* and *vanSB.* The genes *vanSA* and *vanYA* were individually (and uniquely) observed in sample 8 (unassigned ST) and sample 23 (ST203) respectively.

One chromosomal exclusive gene, *aac(6’)-Ii* (aminoglycoside resistance), was observed to be ubiquitous in all samples and one other *msrC* (macrolide and streptogramin B resistance), was observed in all isolates except sample 23 (ST64). The combination of these two genes is reported to confer resistance to compounds from the aminoglycoside, lincosamide, macrolide, oxazolidinone, phenicol, pleuromutilin, streptogramin, and tetracycline classes (129–132). Interestingly, CC94 isolates were not observed to possess any other AMR genes beyond these two examples. The genes *ermB* and *ermT* (conferring resistance to lincosamide, macrolide, and streptogramin) were present in 46 of 68 CC17 isolates. Only one isolate, sample 23 (ST64), was observed to possess both *ermB* and *ermT* genes and each individual gene was distributed relatively unevenly throughout the dataset (*eg. ermB* was observed in 8 ST80 isolates, *ermT* in 5 ST80 isolates, and 5 ST80 isolates lacked both genes); however, it was observed that, if present, an isolate only possessed one of *ermB* or *ermT* on the chromosome (with the exception of Sample 23). The distribution of *erm* in this regard does seem to follow an underlying phylogenetic bias. For example, approximately half of Subcluster δ_2_ possess *ermB* and the other possess *ermT*. This transition appears to have occurred after the divergence of ERR374697 as it is retained in all later diverging taxa (Figures 5,6). Another example of *erm* bias can be observed between Subclusters δ_2_, δ_3_, and δ_4,_ where δ_2_, δ_3_ display bias towards *ermB* and δ_4_ towards *ermT*. As both genes are observed in the Sample 23 chromosome (the earliest diverging δ_4_ taxon), it can be reasonably assumed that this transition occurred after the divergence of Sample 23 from the last common ancestor of the remaining δ_4_ isolates.

The gene *dfrF* (conferring resistance to trimethoprim) was observed distributed throughout the dataset (like *ermBT*) and displayed phylogenetic biases towards Cluster γ, and *ermT* positive members of Subclusters δ_1_, δ_2_, and δ_3_. The gene *tetM* (conferring resistance to tetracycline) was scattered throughout the dataset but was specifically observed in 13 or 18 ST80 isolates.

AMR genes present on either mobilome or chromosome.

Seven of the AMR genes detected were distributed across either the mobilome or the chromosome within the samples investigated (*AAC(6’)-Ie-APH(2’’)-Ia*, *aph(3’)-IIIa*, *ermB*, *SAT*-4, *aad*(6), *dfrG* and *tetL*). This demonstrates the inter-connectedness of the mobilome and the chromosome as a mode of AMR gene transport within these isolates (SI Table 6; Figure 6).

#### Core resistome

Using default pangenomic parameters (106), a core chromosomal resistome (inclusive of soft-core) whereby a gene is represented in ≥95% (≥75 genomes) comprise of two genes: *AAC(6’)-Ii* and *msrC*. When CC94 is excluded, the core mobile resistome (≥64 genomes), is expanded to include *efmA*. A core mobile resistome was not established.

### Single nucleotide point mutations

Linezolid resistance was identified phenotypically in sister taxa, Samples 14 and 24, but no associated plasmid mediated resistance mechanism was identified (SI Table 6; Figure 6). In addition, the 23S rRNA, L3 and L4 gene sequences and amino acid sequences, respectively, were compared with the linezolid susceptible isolates in this study using PointFinder (SI Table 7; Figure 7). No linezolid resistance associated mutations were identified. Thus, the mechanism of linezolid resistance has not been identified. The linezolid resistance phenotype of the BSAC or CC94 isolates was not reported.

The ciprofloxacin resistance phenotype was not reported in the BSAC or CC94 isolate associated metadata. Isolates sequenced for this study were all ciprofloxacin resistant. The gyrase and topoisomerase IV genes (*gyrA* and *parC*) from each isolate was compared with the ciprofloxacin susceptible strains using PointFinder to identify point mutations associated with ciprofloxacin resistance. Mutations in all isolates were identified at amino acid position 83 in GyrA, resulting in either an S83I change in all δ_2_ isolates (Samples 6, 7, 12, 15, 19, 20, and 22) or an S83Y change in the remaining samples sequenced for this study, in *ermT* positive δ_1_ isolates, both ST203 isolates, in the only Cluster α isolate, and in a four-isolate clade of δ_2_. In addition, two mutations at amino acid 709: Y709N and Y709D, outside the quinolone resistance determining region (QRDR) in the same groups of isolates. Point mutations at p.S80 were observed across all CC17 isolates except ERR37474. The majority (53 of 78) of isolates displayed an S80I mutation and the remaining 15 isolates displayed an S80R mutation. Point mutations in ParC follow a phylogenetic pattern, appearing in blocks with sister taxa (Figure) Interestingly, all Cluster γ isolates displayed the S80R mutation. As these point mutations are at the same site, it can be reasonably assumed that this point mutation is more likely to be inherited from the CC17 common ancestor than *via* several individual convergent events.

Thus, ciprofloxacin resistance was due to the mutations at serine 83 in the GyrA protein and serine 80 in the ParC protein in all isolates sequenced for this study.

### Metal resistance

No metal or biocide resistance genes were observed on any mobilome sequence. A “core” metal-resistome of 3 single-copy genes (*chtR, chtS,* and *sodA*) was observed in all chromosomes (SI Table 8; Figure 8). A further 3 single copy genes (*copA*, *copB*, and *copY/tcrY*) observed in all chromosomes except for two isolates sequenced during this study, sample 9 and sample 21 (both from ST192). One gene, *perR* was observed as a single-copy ortholog in all clinical isolates (inclusive of ST192) but absent in all “Cork” samples. One gene, *dpr/dps* was observed exclusively in sample 8 (unassigned ST). Finally, a set of 6 genes (*merA*, *merR1*, *tcrA*, *tcrB*, *tcrY*, and *tcrZ*) were observed exclusively in the “Cork” samples (CC94). In summation, all samples (except ST192) shared 7 genes (*chtR, chtS, copA, copB, copY/tcrY, perR,* and *sodA*). These 7 genes confer resistance towards selenium (*chtR*), hydrogen peroxide (*chtR, perA,* and *sodA*), chlorhexidine (*chtS*), copper (*copA*, *copB*, *copY/tcrY*) and silver (*copB*) (36, 133). The absence of *copABY* from ST192 may indicate copper and silver susceptibility in these samples. The presence of *dpr/dps* in sample 8 is expected to confer iron resistance and increased resistance to hydrogen peroxide (134, 135). The presence of *merA* and *merR1* in the CC94 samples indicates intrinsic resistance to mercury and phenylmercury acetate (an organomercuric compound) and the presence of *tcrABYZ* suggests increased copper resistance (43,136–139).

Using default pangenomic parameters (106), a core chromosomal metal resistome (inclusive of soft-core) whereby a gene is represented in ≥95% genomes (≥75 genomes) comprise of six genes: *chtR, chtS, copA, copB, copY/tcrY* and *sodA.* When CC94 is excluded, the core metal resistome (≥64 genomes), is expanded to include *perR*.

### Virulence factors

Each CC94 isolate contained a plasmid mediated virulence factor (VF), the biofilm-associated transcription factor *bopD* (SI Table 9; Figure 9). Only one other VF was observed to be plasmid mediated: the fibrinogen binding surface protein *fss3* (only observed within sample 18 (ST80)), all other observations were chromosomally mediated. All isolates were observed to possess *clpP* (a caseinolytic protease), *cpsA* (a biofilm associated undecaprenyl diphosphate synthase) and *cpsB* (a capsule associated phosphatidate cytidylyltransferase). A collagen adhesin precursor (*acm*) was observed in all isolates except sample 1 (ST1421) and the cell wall anchor protein *sgrA* was observed in every isolate except sample 26 (ST202). All isolates in CC17 (except Sample 1 (ST1421)) contained the bile salt hydrolase *bsh* and all CC17 isolates except ERR375031 (ST17) contained the endocarditis/biofilm-associated pilus *ebpC.* The collagen binding microbial surface components recognizing adhesive matrix molecule (MSCRAMM) gene *ecbA* was observed in sample 17 (ST80), ERR375043 (ST16), ERR374741 (ST17), ERR374767 (ST64), and in two ST203 samples (ERR374813 and ERR374914). The collagen adhesin protein *scm* was observed in 14 of 16 ST80 isolates, 5 of 17 ST17 isolates, 2 of 5 ST18, 7 of 9 ST78, 1 of 3 ST192 (sample 21) and in sample 26 (ST202). Finally, *fss3* was observed in 14 of 16 ST80 isolates, all 6 ST203 isolates, in both ST132 isolates, in one isolate each from ST16, ST17, ST64, ST78, ST192, ST787, ST1032, and ST1421 and one unclassified ST isolate (sample 22). Both *fss3* and *scm* were absent from sample 17 (ST80).

Using pangenomic parameters, a core virulome (inclusive of soft-core) whereby a gene is represented in ≥95% (≥75 genomes) would yield *acm, bopD, clpP, cpsA, cpsB, efaA, hasC, htpB, lap,* and *sgrA*, illustrating a key pathenogenic potential for biofilm-associated proteins, capsular polysaccharide biosynthesis, caseinolytic protease, endocarditis specific antigen and collagen adhesins (140, 141). When only clinical isolates are considered, the core virulome (≥64 genomes) is expanded to include *bsh, epbA* and *epbC,* further increasing cardiac virulence and limiting the antibacterial function of bile acids (142, 143).

### Phages

Prophages were observed in the chromosome of 65 isolates (2.59±1.02; 1 ≤ *n* ≤ 4) and all were observed to be non-transposable (SI Tables 17-19). The majority (196 of 202) of prophages were predicted to be Siphoviridae, three prophages were observed to be either Siphoviridae or Myoviridae in ERR374794, ERR374824, ERR375046, one prophage (from ERR374806) was observed to be Myoviridae, one prophage (from sample 25) was observed to be Inoviridae, and one prophage (from sample 20) was of unknown taxonomy. Prophages were not observed within any mobilome. One resistance gene, *ermB*, was observed in a prophage of ERR374741, however no other resistance mechanism (antibiotic or metal/biocide) and no virulence factors were observed on any other prophage. Without this *ermB* phage, ERR374741 would be susceptible to lincosamide, macrolide, and streptogramin. The prophage genome within ERR374741 was also observed to contain a probable secondary metabolism mechanism, however the product class could not be determined. One prophage within sample 13, sample 28, and ERR374909 was observed to contain a probable polyketide synthase. When these prophages were searched using Mash (SI Table 19), a set of 4 phages were mapped to 40 of 79 genomes. In the “Cork” samples, *Enterococcus* phage ΦFL2A was ubiquitously observed. A total of 17 prophages were observed within 17 BSAC genomes, *Enterococcus* phage EFP01 was observed in 11 genomes, *Listeria* phage B025 was observed in 4 genomes, and *Corynebacterium* phage BFK20 was observed in 2 genomes (Figure 14)

When isolation dates were considered for the BSAC samples, vancomycin resistance did not appear to be affected by the presence of a prophage as both VSE and VRE were represented (57). Both *Corynebacterium phage* BFK20 prophage-containing genomes were isolated in 2009, *Listeria* phage B025 prophage-containing genomes were isolated between 2002 and 2011, and *Enterococcus* phage EFP01 prophage-containing genomes were isolated between 2001 and 2011. In genomes sequenced for this study, *Corynebacterium* phage BFK20 was observed in 1 genome (sample 7), *Enterococcus* phage EFP01 was observed in two genomes (samples 13 and 28), *Enterococcus* phage ΦFL2A was observed in sample 11, *Listeria* phage B025 was observed in 3 genomes (samples 6, 12, and 19), and a duo of *Enterococcus* phage ΦFL2A and *Listeria* phage B025 was observed in 5 genomes (samples 2, 14, 22, 23, and 24).

A core phage has previously been observed in studies into *Enterococcus faecalis*, where it is implicated in virulence and genome plasticity (144), however, to our knowledge, this trend has not been observed for *E*. *faecium*.

### Pangenomics

As mentioned above, a pangenome was constructed for this dataset each pangenomic category was populated (*n*_genes_; “core pangenome” (99% ≤ *n*_samples_ ≤ 100%), “soft core pangenome” (95% ≤ *n*_samples_ < 99%), “shell pangenome” (15% ≤ *n*_samples_ < 95%), and “cloud pangenome” (*n*_samples_ < 15%)). The number of genes in the “whole genome” core and soft-core pangenomes and the “chromosomal” core and soft-core pangenomes are relatively unaffected by the mobilome (1324 and 236 genes *vs* 1327 and 229 genes respectively (SI Table 12)). However, the shell pangenome (2252 genes *vs* 1550) and the cloud pangenome (3306 genes *vs* 2762 genes) were more pronouncedly affected.

### Pangenomic enrichment

The core pangenome (inclusive of all isolates) was observed to be enriched (*P*_BD_ ≤ 0.005) for major “house-keeping” and viability functions such as metabolism (*eg.* lipid metabolic process (GO:0006629), protein metabolic process (GO:0019538), carbohydrate metabolic process (GO:0005975), and RNA metabolic process (GO:0016070)), transport (*eg.* protein transport (GO:0015031) and ion transport (GO:0043167)), stress response (GO:0006950), signal transduction (GO:0050794), biosynthesis (GO:0009058), cellular regulation (GO:0065007), and localization (GO:0051179) (SI Table 13). The soft-core pangenome was observed to be enriched for localization and for transmembrane transport (GO:0022857). The shell pangenome was enriched for similar processes as the core genome, namely metabolic processes, transport, stress response, and localisation. The shell pangenome was also significantly enriched for small molecule metabolism (GO:0044281) and transposition (GO:0032196). The cloud pangenome was also enriched for metabolic processes, transport, and signal transduction. There was only one significant purification: transposition in the core genome. These results suggest a dynamic and non-specific evolution of the *E. faecium* pangenome and provides an insight into their genomic organisation and architecture.

### Co-evolution

When vancomycin resistance genes were sampled from the 28 genome dataset, a significant association (*P* ≤ 0.005) was observed between *vanAHX* and each of *cadC, dinB, gmuD, gmuR, hin, hisB, nag3, sdhB,* and *uxaC* and a significant dissociation with each of *clpC, czcD, czrA, dkgB, dps, entP, fetA, fetB, fixK, fruA, gatA, lgt, mco, metB, opuCA, panE, phoP, rihB,* and *uvrA* (SI Table 14). These results suggest that VR*Efm* carrying *vanA* typically display an increase in carbohydrate processing (*via gmuD, gmuR, nag3, sdhB*, and *uxaC*), cadmium transport (*via cadC*), and DNA repair (*via dinB*) (145–152). Conversely, isolates with a *vanA*+ genotype were also more likely to display genotypes for a decrease in biofilm formation (*via clpC*), lipoprotein synthesis (*via lgt*), copper, iron, and zinc resistance (*via czrA fetAB,* and *mco*), pantothenate biosynthesis (*via panE*), quaternary ammonium compound resistance (*via opuCA*), oxidative resistance (*via fetAB* and *opuCA*), and stress response (*clpC, dps,* and *uvrA*) (153–162). Comparatively, *vanBWXBYB* was observed to be significantly associated with each of *agaS, bfrA, dgaR, dgoA, dgoD, epsF, fruA, gatB, gmuE, gmuR, gpr, Int-Tn, lacD2, manR, manX, metK, mhpE, mro, mshA, noxE, sorC, ssbA,* and *treP* and dissociated with each of *aes, araQ, cysM, dctM, dppC, dppE, oppB,* and *tuf.* These observations suggest that *vanB*+ VR*Efm*, like *vanA*+ VR*Efm* possess increases in carbohydrate (specifically sugar) catabolic processing genotypes (*via agaS, dgoAD, lacD2, manRX, mshA,* and *treB*), however, an increase was observed in biofilm formation genotype (*via epsF*) and stress response (*via treB*) (163–169). Isolates with a *vanB* phenotype were observed to be decreased in phenotypes for arabinose uptake (*via araQ*), cysteine synthesis (*via cysM*), dicarboxylate transport (*via dctM*) and protein transport (*via dppCE* and *oppB*) (170–174). These protein transport genes are essential for sporulation in other Firmicute species, however, as *Enterococcus* are non-spore forming, their exact role with relation to *vanB* has not been elucidated (174–176).

Interestingly, fructose processing genotypes (*via fruA, gatAB,* and *gpr*) are significantly decreased in *vanA*+ isolates while being significantly increased in *vanB*+ isolates (SI Table 16), suggesting a possible fructose metabolic niche in *vanB+* isolates.

### Rampant recombination

Whole genome alignments and subsequent recombination analyses indicate massive recombination events in across the CC17 genome during its divergence from non CC17 *E. faecium* ancestors (Figure 13; SI Figure 3). The majority of recombination appears to have occurred approximately within the first 50% of the genome.

#### Integron evolution

A total of 31 CC17 isolates were observed to possess an intrgron element, where three isolates (Samples 7, 15, and 22 (all Subcluster δ_3_)) were observed to possess two (SI Table 20; Figure 15). Integrons followed a phylogenetic distribution in Subclusters δ_3_ and δ_4,_ however all other clusters were observed to be pseudorandomly distributed throughout the phylogeny. Genes-of-interest (resistance or virulence genes) were only observed on integron containing contigs in Subclusters δ_3_ and δ_4_. All isolates with a gene-of-interest possessed *lsaA* (conferring resistance to clindamycin, quinupristin-dalfopristin, and dalfopristin). Subcluster δ_4_ isolates only possessed *lsaA*. All Subcluster δ_3_ isolates also possessed *efmA* (an efflux pump conferring resistance to macrolides and fluoroquinolones), *chtS* (conferring resistance to chlorhexidine) and *sodA* (conferring resistance to peroxides). A clade within Subcluster δ_3_ (comprising of Samples 6, 12, 19, and 22) also all contained *efaA* (an endocarditis specific antigen) on their integron containing contigs.

### Observation of an unusual contig

Five isolates (Samples 2, 14, 24, and 25 (all assigned to ST80; subcluster δ_4_) and Sample 18 (ST80; Cluster β)) possessed a contig with a copy of *arnB,* a UDP-4-amino-4-deoxy-L-arabinose--oxoglutarate aminotransferase involved in Gram-negative Lipid A biosynthesis and polymyxin resistance (177–179). The contig in samples 2, 14, and 24 are identical (SI Figures 4(a)-4(c)), however those attributed to samples 18 and 25 are distinct (SI Figures 4(d)-4(e)). Samples 2 and 24 were isolated from the same patient and all three of these samples are phylogenetic neighbours (Figure 5). These contigs have a plethora of cell wall synthesis and lipopolysaccharide modification genes which may play a role in resistance. Of particular interest, sample 17 contained phage sequences suggesting a possible viral mediated horizontal gene transfer. Instances of chromosomal segment transfer between Gram-positive and Gram-negative species (and *vice-versa*) has previously been attributed to phage activity and plasmid integration (180–182).

## Discussion

Within Ireland VRE has already spread nationally and is endemic to Irish hospitals (183). The level of VRE in blood stream infections in Ireland has remained at or above 30% since 2005. While some other EU countries have levels above 30%, they have not consistently had this level of VRE BSI over such a wide timeframe. In 2019, >25% of VR*Efm* BSI in eleven additional EU countries have been identified using the ECDC surveillance data. The spread and dominance of VR*Efm* within Irish hospital patients over such a long duration provides us with a unique setting to study the changing dynamics and evolution of VR*Efm*. A recent study of the global dissemination of *E. faecium* identified that it has two main modes of genomic evolution: the acquisition and loss of genes, including antimicrobial resistance genes, through mobile genetic elements including plasmids, and homologous recombination of the chromosome. Within this global study 261 genomes contained the *vanA* gene. Within the hospital associated A1 clade of *E. faecium* the median number of AMR genes increased in the global study between 2001 and 2019 from 8 to 11 in 2015. Within this study for the *Efm* isolated between 2001 and 2011 the total AMR gene numbers corresponded with the presence of *vanA* rather than year. The vancomycin susceptible isolates and those with the *vanB* gene had less AMR genes than those with the *vanA* gene. *vanA*+ isolates had between 13 and 20 AMR genes, while those with *vanB* or no van gene had a total of between 5 and 12 AMR genes (except one vancomycin resistant isolate with *vanB* (*n* = 15) and one van susceptible isolate with no van genes (*n* = 16)). The timeframes for both groups were distributed across the ten years. Those with the highest AMR gene numbers both contained *vanA* and *vanB*. The number of AMR genes from the *vanA* study in 2018/2019 was between 12 and 18, which is consistent with the number from the previous study. Thus, the number of AMR genes in Ireland did not increase over time.

The aims of this study were to compare the pan-genomes, mobilomes and chromosomes of the available genome sequences of vancomycin resistant or susceptible *E. faecium* across the timeframe that VRE (analysed by the ECDC (2002 to 2019)) increased in prevalence from 11.1% to 38.4% in Ireland. To monitor and limit the spread of VR*Efm* we need to understand how it evolves and acquires vancomycin resistance, how transmission networks are operating and how VR*Efm* is developing resistance to last-line antibiotics.

The same AMR gene was not observed in both chromosome and mobilome in any instance, which indicated genomic control of genetic redundancy (184–186). Vancomycin resistance was observed to be either plasmid mediated (*via vanA*) or chromosomally mediated (*via vanB*), and rarely a combination of *vanB* on the chromosome and plasmid mediated *vanA* vancomycin resistance was observed. Tetracycline (and probable tigecycline) resistance (*via tet*(*L*)) was differentially dispersed in clinical isolates, with *tet*(*L*) being observed in the mobilomes of ST17, ST18, and ST203 and in the chromosomes of ST80 (if present within the genome). Other tetracycline resistance genes implicated in tigecycline resistance, *tet*(*M*) and *tet*(*U*), were observed in addition to *tet*(*L*) on the chromosome of most ST80 isolates. Interestingly, in contrast to previously reported trends (187, 188), increased metal resistance did not positively correlate with increased drug resistance, in fact the opposite was observed, with the “Cork” strains displaying diminished drug resistance and increased metal resistance.

Core-genome MLST corroborated phylogenetic and hierarchical clustering results, suggesting that core populations of *E. faecium* persist in Irish hospitals with frequent evolution towards pathogenicity *via* plasmid mediated transfer of *vanA* or chromosomal incorporation of *vanB* (Figures 5,12). This suggestion is corroborated by the genomic similarity between isolates in different clusters and clades (regardless of isolation year) and by the fact the VSE is observed within Cluster β and clades δ_1_ and δ_4_ (Figure 5).

We determined that a relatively stable “prophagome” exists between isolates (SI Table 17), however, these are not, to our knowledge, implicated in the evolution of resistance or pathogenicity, as observed with closely related species (189, 190).

An interesting potential virulence mechanism difference was observed between *vanA*+ and *vanB*+ isolates, whereby *vanA*+ genotypes were unlikely to display a biofilm forming genotype (*via clpC*) and *vanB*+ isolates were likely to display an *epsF* biofilm forming genotype (SI Table 16). Both vancomycin resistance genotypes displayed increases in carbohydrate catabolism genotypes, however *vanB*+ isolates were more likely to display specific sugar processing genotypes. Interestingly, *vanA*+ genotypes were less likely to coevolve with metal resistance, oxidative stress resistance, panthenoate, or lipoprotein synthesis genotypes. These results suggest that *vanB*+ isolates are more likely to have evolved due to different stressors than *vanA+* isolates. As *vanA* alignments used in this manuscript did not have any point mutations (regardless of the year of isolation), it is likely that a narrow range of mobile genetic elements confer resistance to commensal VSE in the immunocompromised, prompting their pathogenicity.

The significant differences in observed gene density (higher in mobilomes), GC% (lower in mobilomes), gene length (shorter in mobilomes) coupled with insignificant in coding proportions between chromosomes and mobilomes suggest that mobilomes are more likely to evolve either through the acquisition of statistically shorter sequences (while maintaining an equivalent coding potential) or through successive gene fission and/or gene subfunctionalisation events (191–194). While not statistically significant in most samples, coding nucleotide density is, on average, approximately 7% less in mobilomes (SI Tables 11-12), where significantly variant coding proportions are also observed (σ_2chromosomal_ = 1.04; _mobilomal_ = 3.62; *P* = 1.99*e* (Levene’s test)). It is possible that these variances may just be an artifact of the highly variant sequences distributed across plasmids when compared to more stable chromosomes, these variances may also underly nucleotide deletion “clean up” processes following subfunctionalisation or fission events indicating evolution towards a more streamline, and therefore less bioeconomic expensive plasmids, thus promoting their propagation (193,195–197).

The pangenome of *E. faecium* is relatively open, with core and soft-core components only accounting for approximately 22% of the pangenome (when the mobilome is included) and for 26.5% when the mobilome is excluded (SI Table 12). This variation, coupled with the non-specific functional enrichments observed within the pangenome (SI Table 13), highlights the rampant evolutionary capabilities in “fixed” genomic biomolecules. This variation may be partially explained by the rampant recombination observed throughout CC17 or the diversity of prophages observed throughout the respective genomes, however more research is needed to confirm the role of phages on VRE genome (Figures 11-13; SI Figure 3). As observed for phage data, integrons were observed throughout CC17 (SI Table 19; Figure 15). When integrons were observed in phylogenetic blocks, genes-of-interest were observed (SI Table 20; Figure 15). However, as these genes are widely observed in *E. faecium*, it is possible that these integrons are faciliting mobilisation of these genes. While recombination has been reported in *E. faecium* (198, 199), the extent of recombination observed during this study has not, to our knowledge, been previously reported. This recombination may have been a driving force in the evolution of *E. faecium* CC17 towards pathogenicity (200). Despite the rampant variation observed between genomes used in this study, chromosomal size, gene content, gene density, and the phylogenetic proximity of each genome remained largely stable throughout the dataset. The relatedness of the VSE and VRE indicates that vancomycin susceptible *Efm* are transferring between patients and then acquiring plasmids to become VR*Efm* rather than or in addition to the movement of VRE between patients. Stability of *Efm* within Ireland over the 20 years indicates that the increases in VRE across patients is not due to frequent and multiple introductions of different *Efm* into the hospital but rather the maintenance of genomic-related susceptible *Efm* that acquire vancomycin resistance plasmids and the spread of a cohort of highly related VR*Efm*. Thus, the susceptible and resistant *Efm* are highly related and fixed within the hospital patients. To tackle VRE within Ireland we need to focus on reducing VSE in addition to reducing VR*Efm.* This requires analysis of the epidemiology of VSE, VR*Efm* and the plasmids containing the *vanA* gene. While this study provides some understanding of the past, we need to perform studies on the VS*Efm* and VR*Efm* within our hospitals to understand the continuing rise of VR*Efm* and the infection control required to limit the spread.

## Conclusion

We have demonstrated that the genomic characteristics of the studies VS*Efm* and VR*Efm* (such as genome size, gene density, and gene count) are relatively stable across space and time despite an open and plastic pangenome. The relatedness of the VS*Efm* and VR*Efm* across time indicates that to reduce or remove VR*Efm* from Irish hospitals we must concurrently remove VS*Efm*. The problem of vancomycin resistance within these pathogens is plasmid mediated *vanA* rather than chromosomally mediated *vanB*.

## Author contributions

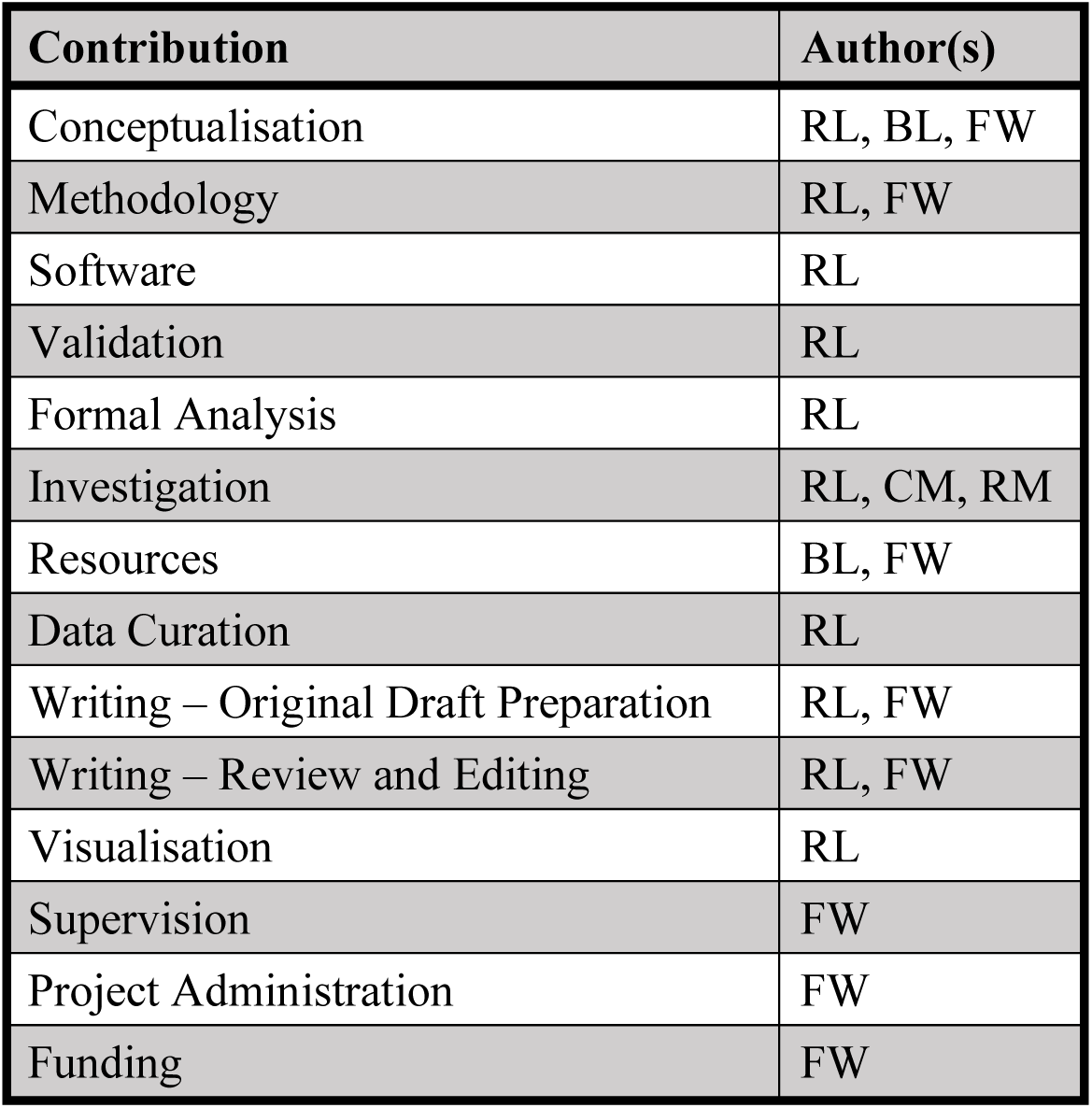

## Acknowledgements

We wish to thank the staff in the microbiology laboratories at MMUH and the BSAC surveillance team for making their data available.

---

**SI Figure 1:**
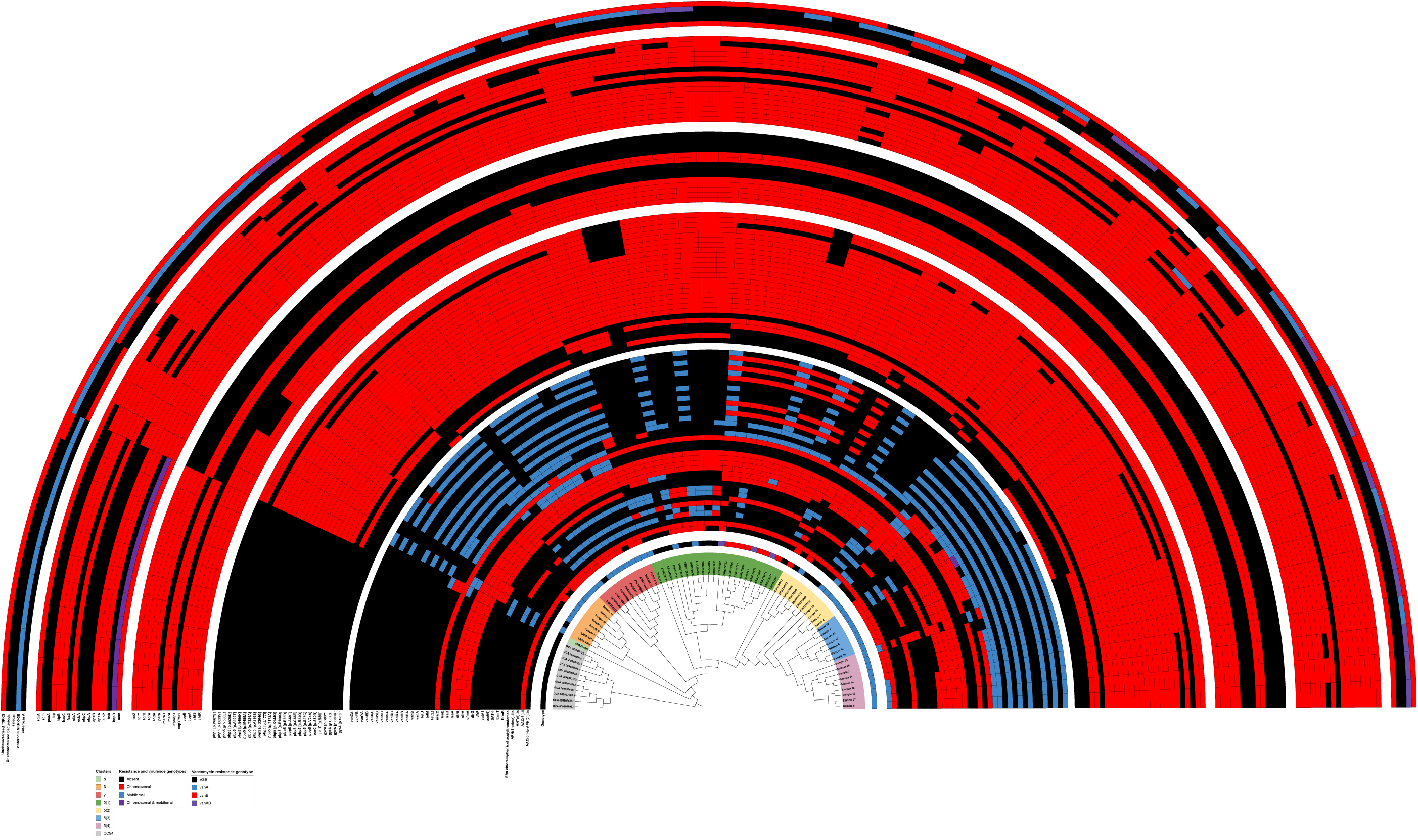
A 10,000 bootstrap phylogeny with all clinically relevant genes arranged as semi-circular categorical heatmaps. Annotation semi- circles are arranged as follows (innermost to outermost): drug resistance genotype, point mutation drug resistance genotype, metal (biocide) resistance genotype, virulence factor genotype, secondary metabolism production genotype

**SI Figure 2:**
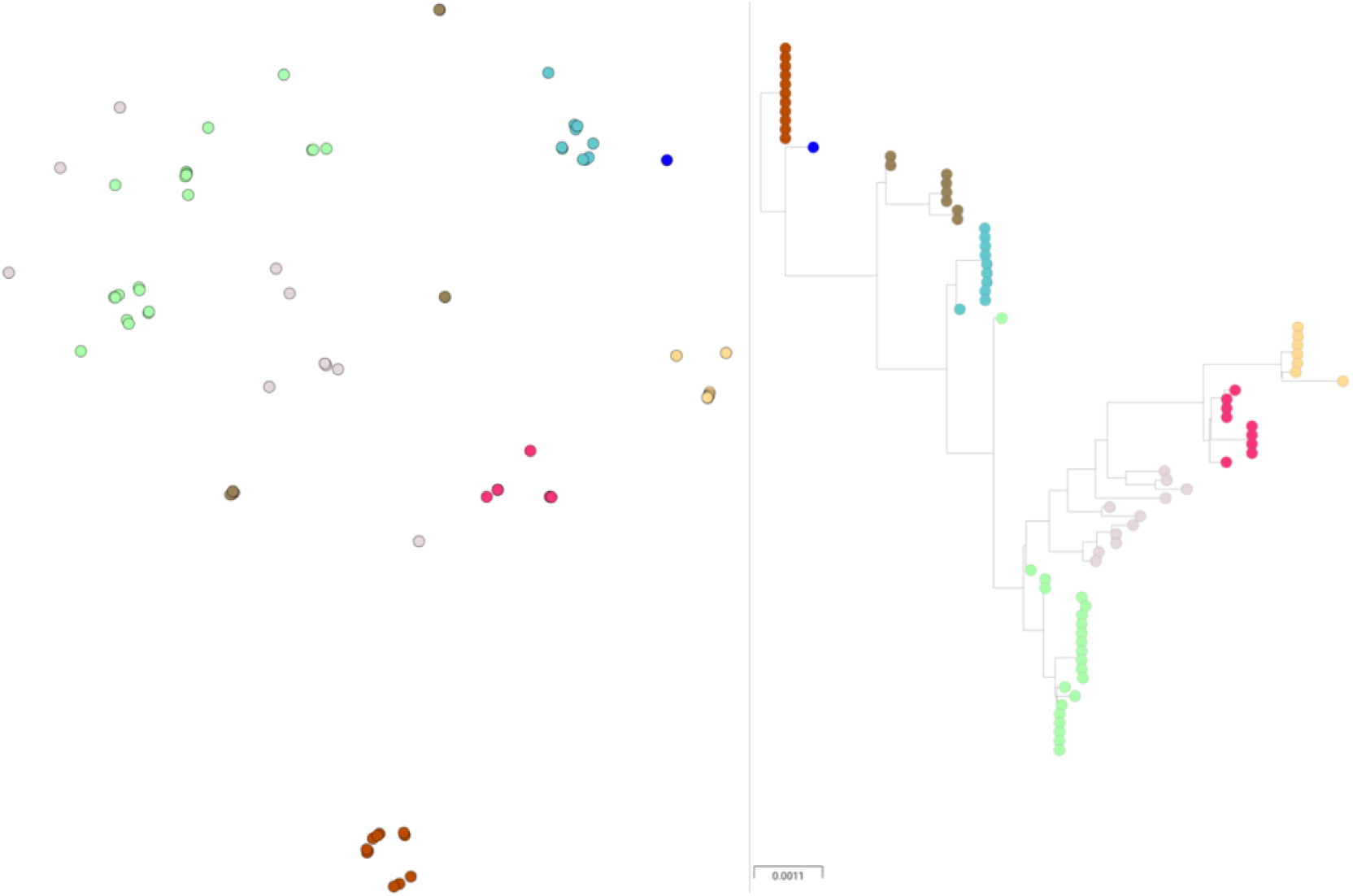
Core genome clustering. These clusters were produced using PANINI to corroborate cgMLST data, the phylogeny on the right is the same phylogeny used in all other images, colours are RheirBAPS clusters

**SI Figure 3:**
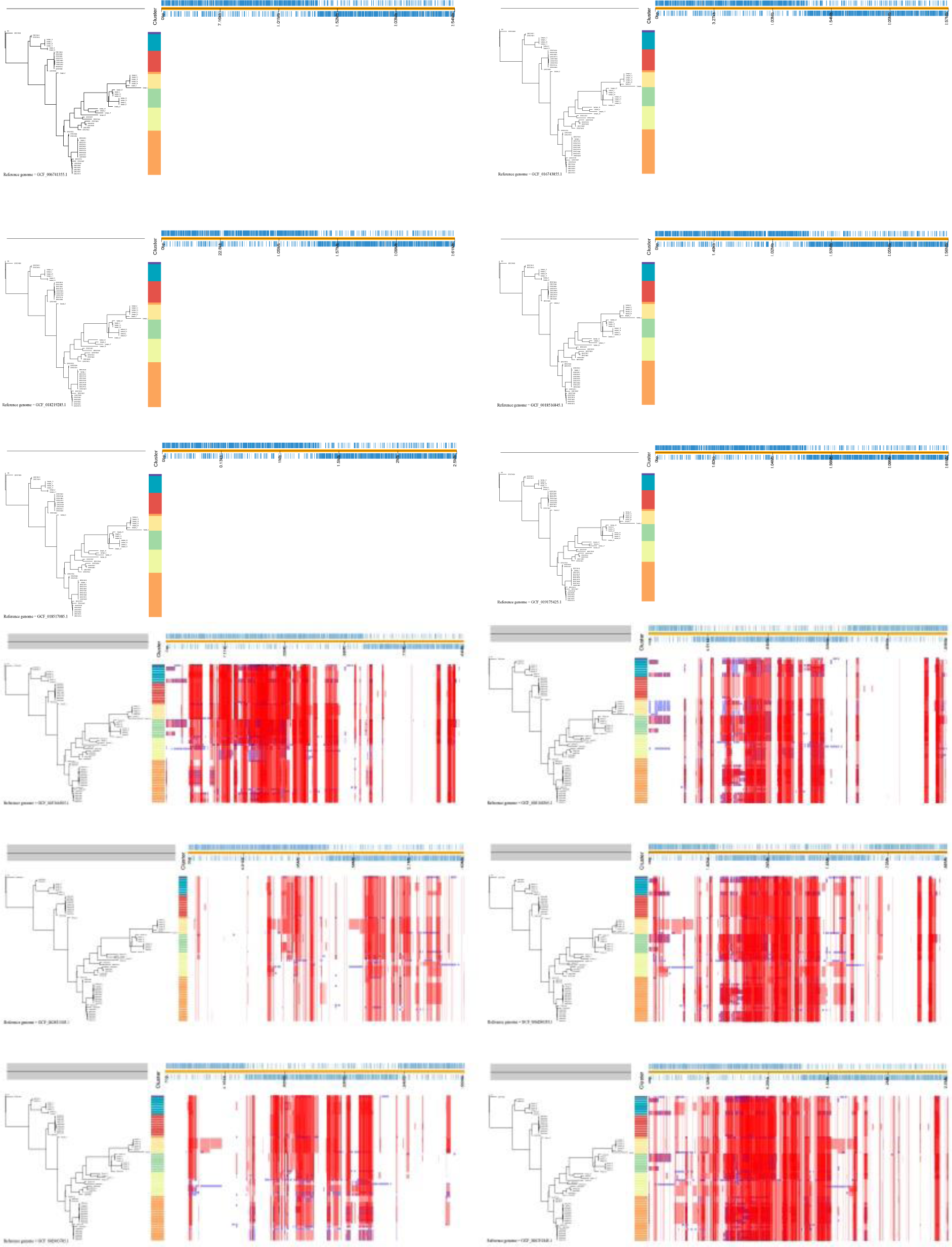
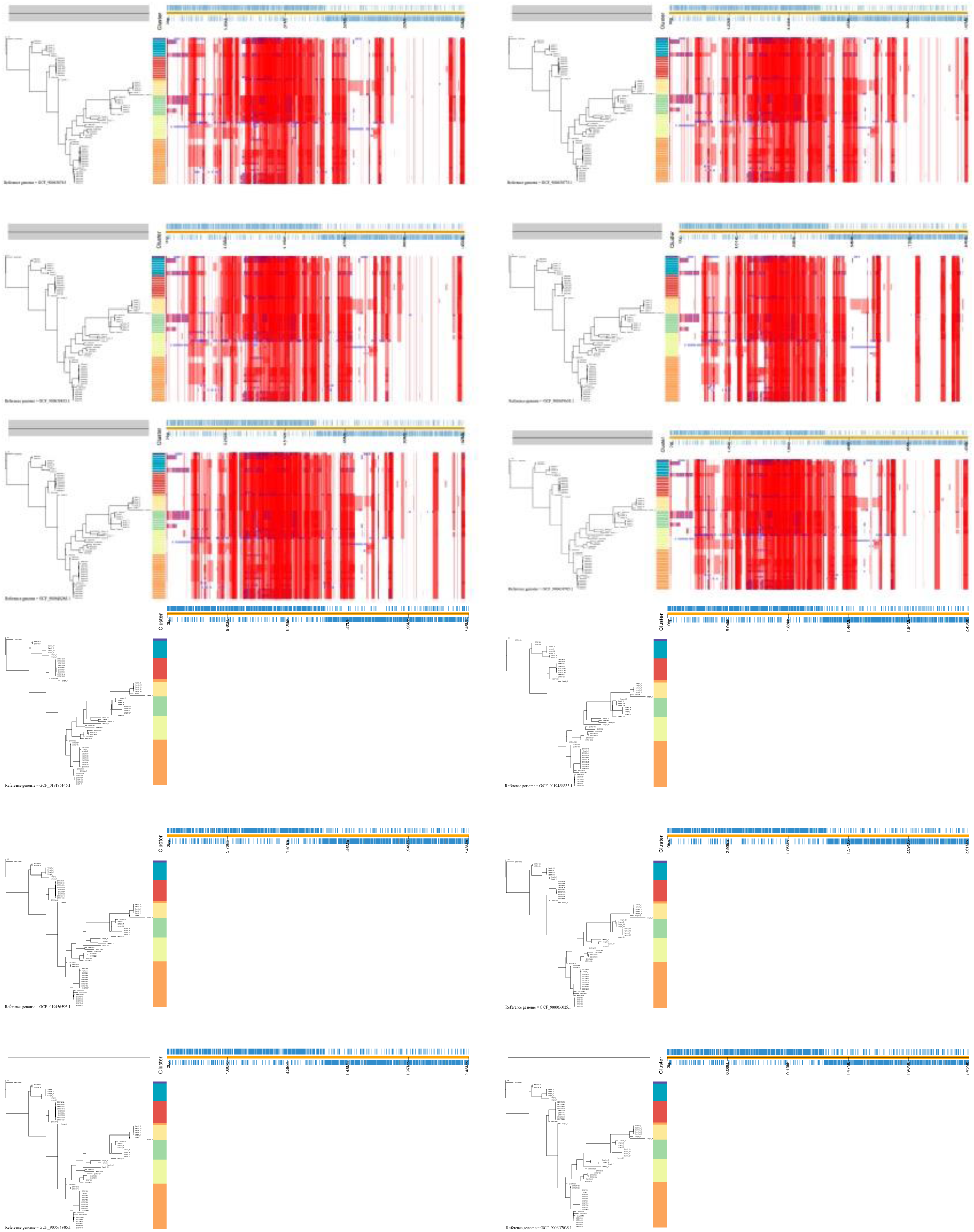
Genome recombination between 24 different reference genomes. Each reference genome diverged prior to CC17. Red bars illustrate hot spots compared to a reference, where a hotspot is defined as a site where more than one species reports a ≥50% likelihood for the same recombination event. Blue boxes illustrate a recombination event for a single species. Multiple different recombination events can occur at the same site (appearing as overlap in this image). The plot at the bottom shows the maximal amount of species affected by a detected recombination event in a given hotspot. Each reference genome is given in the bottom left.

**Figure 4(a):**
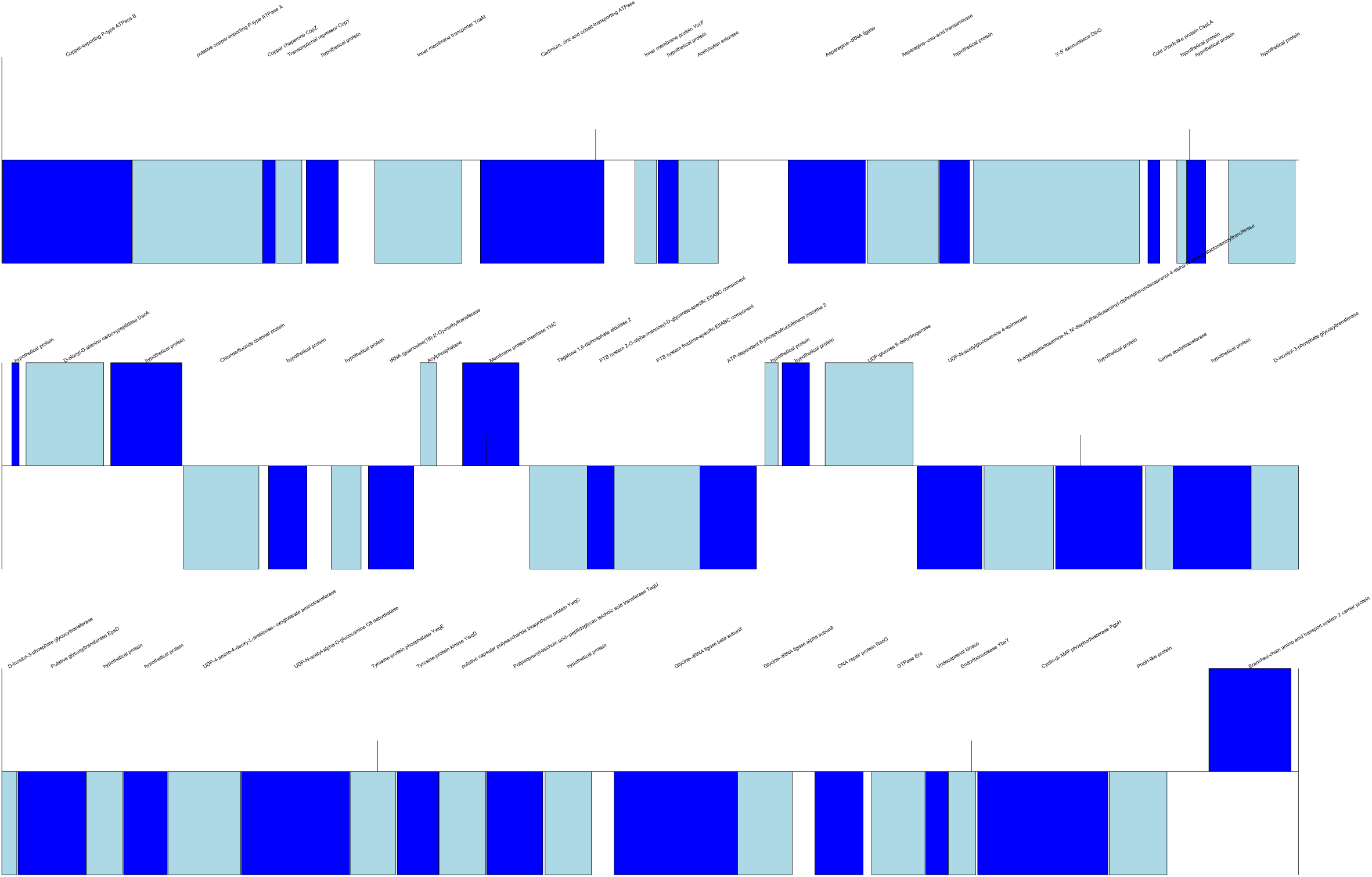
*arnB* contig in Sample 2. Positive sense genes are given above the line and negative sense are below the line. Genes are coloured in alternating blue for ease of visualisation

**Figure 4(b):**
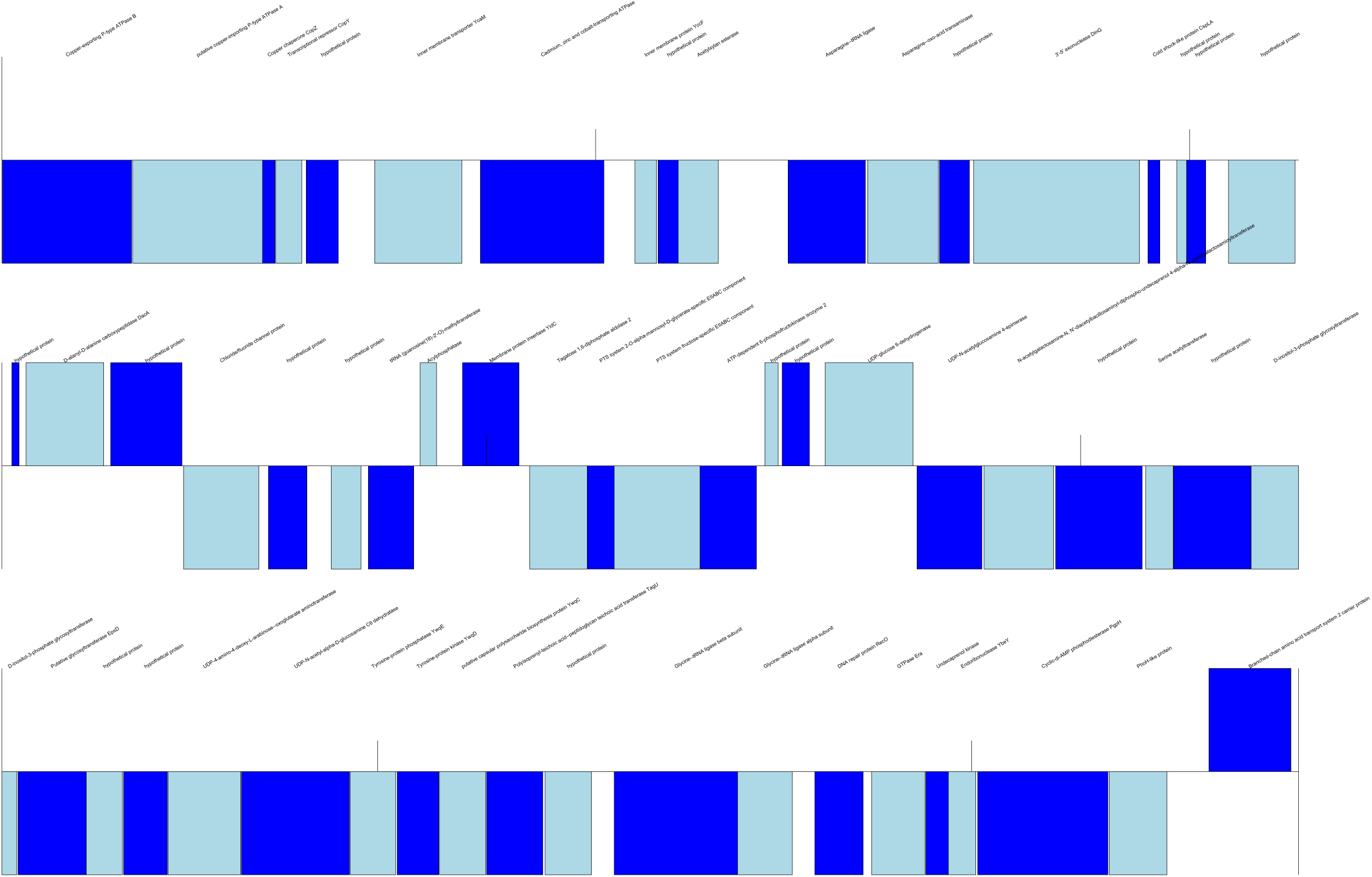
*arnB* contig in Sample 14. Positive sense genes are given above the line and negative sense are below the line. Genes are coloured in alternating blue for ease of visualisation

**Figure 4(c):**
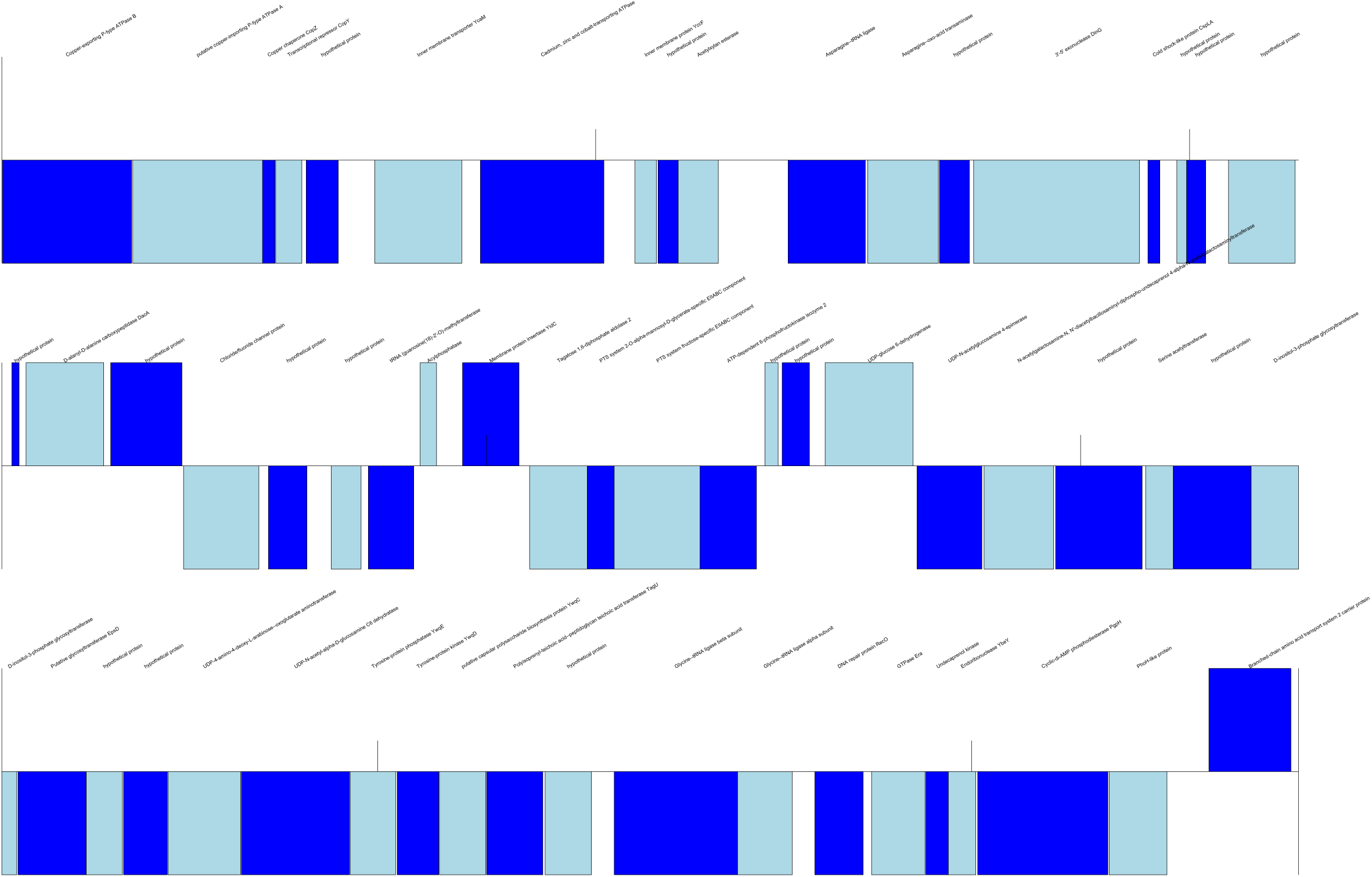
*arnB* contig in Sample 24. Positive sense genes are given above the line and negative sense are below the line. Genes are coloured in alternating blue for ease of visualisation

**Figure 4(d):**
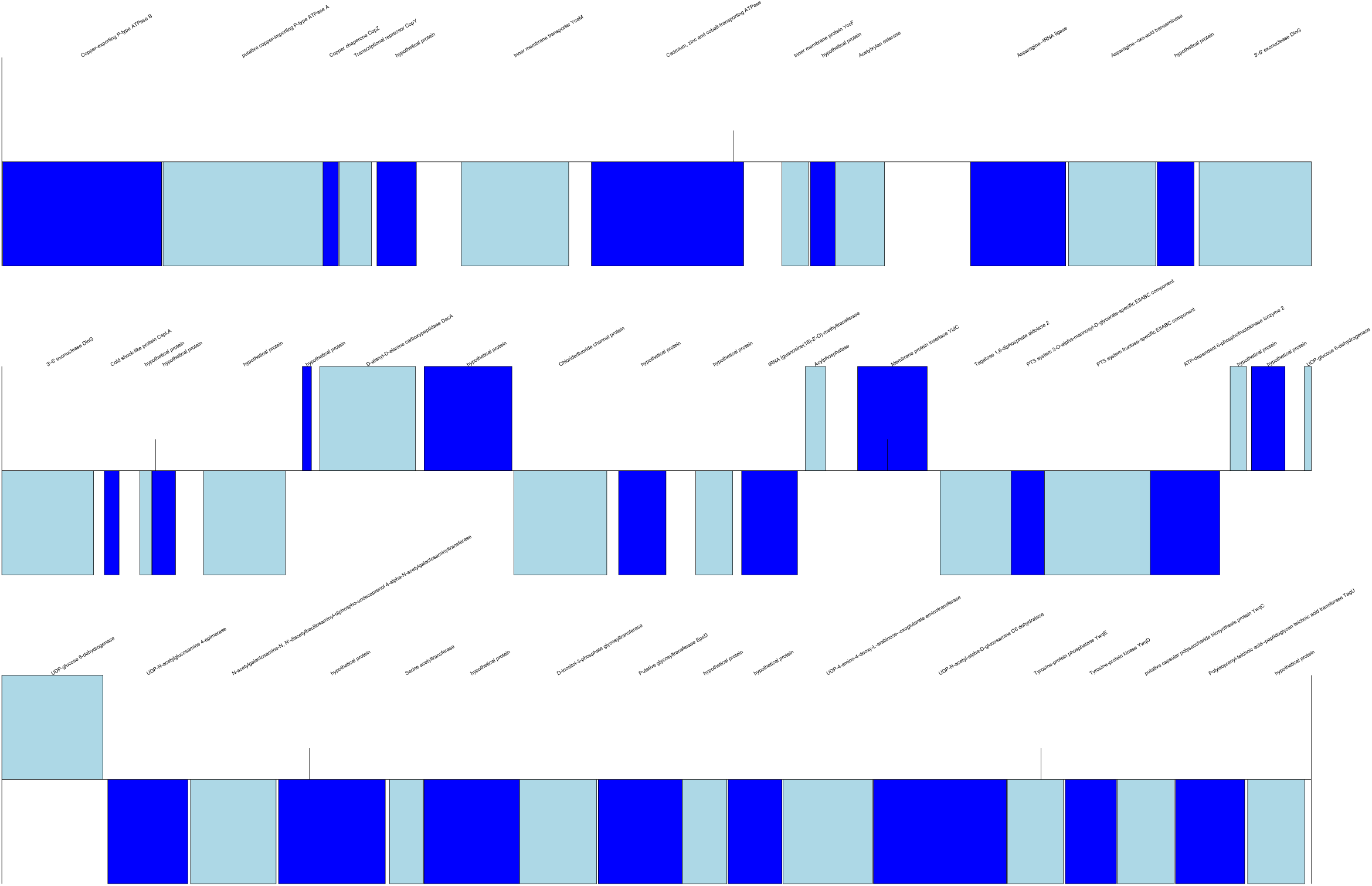
*arnB* contig in Sample 18. Positive sense genes are given above the line and negative sense are below the line. Genes are coloured in alternating blue for ease of visualisation

**Figure 4(e):**
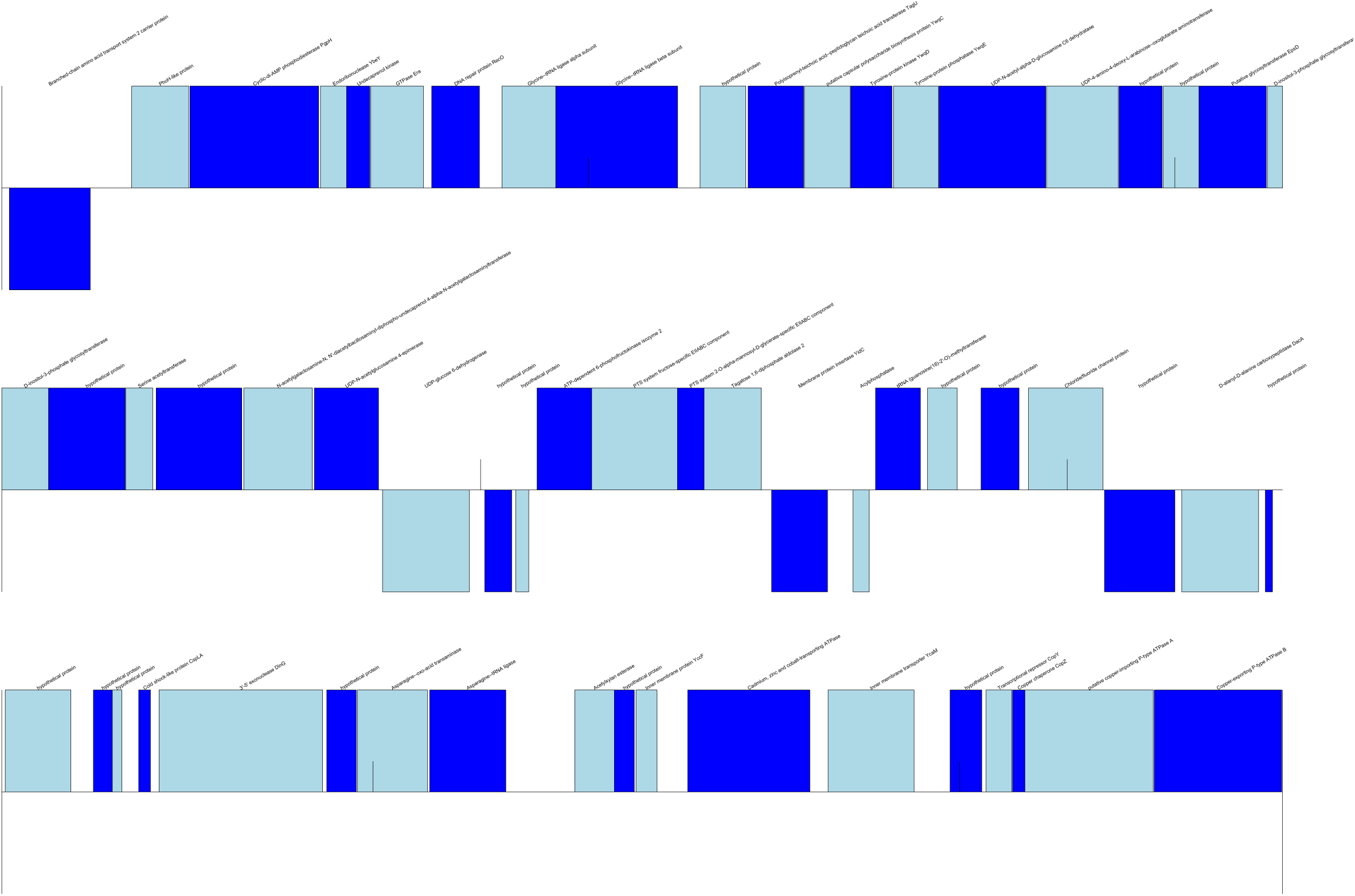
*arnB* contig in Sample 25. Positive sense genes are given above the line and negative sense are below the line. Genes are coloured in alternating blue for ease of visualisation

